# RBM20 phosphorylation on serine/arginine domain is crucial to regulate pre-mRNA splicing and protein shuttling in the heart

**DOI:** 10.1101/2020.09.15.297002

**Authors:** Mingming Sun, Yutong Jin, Chaoqun Zhu, Yanghai Zhang, Martin Liss, Michael Gotthardt, Jun Ren, Ying Ge, Wei Guo

## Abstract

Molecular and cellular mechanisms of mutations of splicing factors in heart function are not well understood. The splicing of precursor mRNA is dependent on an essential group of splicing factors containing serine-arginine (SR) domain(s) that are critical for protein-RNA and protein-protein interaction in the spliceosome assembly. Phosphorylation of SR domains plays a key role in splicing control and the distribution of splicing factors in the cell. RNA binding motif 20 (RBM20) is a splicing factor predominantly expressed in muscle tissues with the highest expression level in the heart. However, its phosphorylation status is completely unknown up-to-date. In this study, we identified sixteen amino acid residues that are phosphorylated by middle-down mass spectrometry. Four of them are located in the SR domain, and two out of the four residues, S638 and S640, play an essential role in splicing control and facilitate RBM20 shuttling from the nucleus to the cytoplasm. Re-localization of RBM20 promotes protein aggregation in the cytoplasm. We have also verified that SR-protein kinases (SRPKs), cdc2-like kinases (CLKs) and protein kinase B (PKB or AKT) phosphorylate S638 and S640. Mutations of S638A and S640G reduce RBM20 phosphorylation and disrupt the splicing. Taken together, we determine the phosphorylation status of RBM20 and provide the first evidence that phosphorylation on SR domain is crucial for pre-mRNA splicing and protein trafficking. Our findings reveal a new role of RBM20 via protein shuttling in cardiac function.

## Introduction

Heart failure (HF) is a major public health issue^1^. It remains the leading cause of morbidity, mortality, and hospitalization among adults and the elderly^2-3^. Given these clinical observations, new therapeutic approaches are urgently needed, which requires the in depth understanding of the molecular mechanisms responsible for the development and progression of HF to develop new therapeutic targets. Posttranscriptional regulation of RNA binding proteins (RBPs) in heart function has not been well studied but is promising study area to develop new therapeutic targets for HF. RNA binding motif 20 (RBM20) is found to be a muscle specific RBP and its genetic defects are linked to dilated cardiomyopathy in human and animal models^4-13^. In a rat model, thirteen out of fourteen *RBM20* exons (exons 2 to14) were deleted and the titin gene (*TTN*) was translated to the largest isoform due to the alternative splicing regulated by RBM20^4-6, 14-15^. The role of RBM20 as a primary regulator of titin pre-mRNA splicing was also verified through a gene-edited mouse model^16^. In this mouse model, the RNA recognition motif (RRM) of RBM20 was depleted, resulting in the expression of the largest titin isoform in the heart of these mice^16^. In addition to titin pre-mRNA splicing, at least thirty other genes such as myosin heavy chain 6 (Myh6), ryanodine receptor 2 (Ryr2), calcium/calmodulin-dependent protein kinase type II d (Camk2d) have also been found to be regulated by RBM20^6, 14, 17-19^. Aberrant splicing through RBM20 depletion in animal models and RBM20 mutations in human patients is also associated with dilated cardiomyopathy (DCM) and subsequent heart failure development^7-13^. Uncovering the splicing mechanism regulated by RBM20 becomes critically important for seeking the treatment of cardiac dysfunction.

RBM20 protein contains two typical domains, RRM domain located at the N-terminus and the serine-arginine (SR) rich domain at C-terminus, which are also commonly found in other splicing factors^20-22^. Both RRM and SR domains play an essential role in defining the splicing sites and the selection of exon/intron boundaries^20, 23^. Multiple serine residues in SR domain can be phosphorylated by multiple kinases including SR-protein kinases (SRPKs), cdc2-like kinases (CLKs) and protein kinase B (PKB or AKT) kinases. These kinases modulate key cellular signaling pathways such as the PI3K/Akt signaling pathway^24-26^. Phosphorylation on SR domain has impacts on localization and activity of splicing factors. Both hyper- and hypo-phosphorylation of splicing factors can suppress splicing reactions^27^ and translocation between the nucleus and the cytoplasm^28-29^. Although SR protein dephosphorylation is essential for the maturation of splicing catalysis and the spliceosome^30^, dephosphorylation has also been found in the export of splicing factors and processed mRNA to the cytoplasm for protein translation^31^. Hence, SR domain phosphorylation and dephosphorylation are critical for numerous steps in splicing factor subcellular localization and splicing control.

However, the phosphorylation status of RBM20 and its role in splicing control and subcellular localization have not been investigated yet. In the present study, we determined the post-translational modifications (PTMs) of RBM20 by middle-down mass spectrometry (MS)^32-33.^ We also investigated the influences of RBM20 PTMs on splicing control and protein shuttling. Particularly, we studied the role of phosphorylation on SR domain in these biological events.

## Method and Materials

### Experimental animals and sample preparation

This study was performed with rat crosses of Sprague-Dawley (SD) x Brown Norway (BN) (All strains were originally obtained from Harlan Sprague Dawley, Indianapolis, IN). Animals were maintained on standard rodent chow. This study was carried out in strict accordance with the recommendations in the Guide for the Care and Use of Laboratory Animals of the National Institutes of Health. The procedure was approved by the Institutional Animal Use and Care Committee of the University of Wyoming. Primary neonatal cardiomyocytes were isolated and cultured from one-day old rats. Mouse heart tissues were also used in this study. Heart tissues from constitutively activated AKT mice mimicking the overexpression of AKT (overexpression mouse model, 2-month old) and AKT2 knockout (KO) mouse models (2-month old) were given as generous gifts from Dr. Jun Ren’s Lab (University of Wyoming). All samples for protein gel electrophoresis were prepared as described previously^14, 34^.

### Primary cultures of neonatal rat cardiomyocytes (NRCMs) and sample preparation

Primary cultures of NRCMs were prepared from one-day old rats using the neonatal cardiomyocyte isolation system as described previously^35-36^. The cells were re-suspended in complete medium (M199/DMEM media and 20% fetal calf serum (FCS) supplemented with 1% penicillin/streptomycin), plated at a density of 1⨯10^6^ cells per cm^2^, and maintained in 5% CO_2_ at 37 °C. Cells were cultured for two days in complete media, and then supplemented with SRPK1 inhibitor SRPIN340 (25 μM) (Sigma-Aldrich, St. Louis, MO, Cat#SML1088) and CLK1 inhibitor TG003 (10 μM) (Sigma-Aldrich, St. Louis, MO, Cat#T5575) respectively for 0 min, 5 min, 1h, 6h, 12h, 24h, and 48h. The vehicle DMSO treatment was used as control. Harvested cells were lysed for protein and RNA preparation.

### Plasmid constructs

Plasmids of pEGFP-C1-8his-RBM20 WT/mutations, pcDNA3.1-mcherry-SRPK1, and pcDNA3.1-mcherry-CLK1 were constructed by Gene Universal, Inc., USA. pcDNA3-Myr-HA-Akt2 was a gift from William Sellers (Addgene plasmid # 9016). pcDNA3.1-E64-70 expressing titin exons 64-70 was constructed as described previously^15^.

### A double reporter splicing assay in mammalian cell system

Human embryonic kidney (HEK29) cells (Life Technologies GmbH, Darmstadt, Germany) were maintained in high glucose Dulbecco’s modified eagle medium (DMEM) supplemented with 10% fetal bovine serum (FBS) (Sigma-Aldrich Chemie GmbH, Munich, Germany [F7524, Lot#124M3337]) and 1% penicillin/streptomycin 10,000 U/mL (Gibco by Life Technologies GmbH, Darmstadt, Germany). For transfection, 25,000 cells/well were seeded on 96-well Nunc F96 MicroWell™ plates (Life Technologies GmbH, Darmstadt, Germany) and transfected with a total of 200 ng of plasmid DNA of which 1 ng was splice reporter (PEVK Ex4-13) plus a corresponding amount of RBM20 WT, RBM20 mutations or control plasmid (pcDNA3.1) in a 20x molar excess. To deliver plasmid DNA, we used the 40 kDa linear polyethylenimine (Polysciences Europe GmbH, Hirschberg, Germany) at a 1:3 ratio (DNA: PEI40). Cells transfected were at a confluence of 50-60%. Compounds were applied 24 h post-transfection at a final DMSO concentration of 1%. Luciferase activity was measured 60 h post-transfection using the Dual-Luciferase® Reporter Assay System (Promega GmbH, Mannheim, Germany) on an Infinite® M200 Pro (TECAN, Maennedorf, Switzerland) plate reader. Ratios of firefly to renilla luciferase activity were normalized to the control (pcDNA3.1) without the splicing factor. Cell viability was measured 60h post-transfection using a resazurin based staining of metabolically active cells (PrestoBlue®, Life Technologies GmbH, Darmstadt, Germany).

### RT-PCR analysis

Hela cells were grown in DMEM medium supplemented with 10% FBS. TTN mini-gene (Ex64-70) was co-transfected with RBM20 WT or mutated plasmids in Hela cells respectively using Lipofectamine 2000 (Invitrogen, Carlsbad, CA) according to the manufacturer’s instructions. Cells were harvested for protein and RNA preparation at 48 h post co-transfection. RT-PCR was performed to detect exon inclusion or exclusion of TTN mini-gene under each experimental condition using forward primer (5’-ACCAGCTGTGCACACAAAGA-3’) and reverse primer (5’-TCTTCTTTGCCACAGGAACG-3’). Results were confirmed in three independent experiments, and the PCR products were analyzed on ethidium bromide agarose gels as described previously^15^.

### Middle-down mass spectrometry

RBM20 was constructed and expressed with BaculoDirect™ Baculovirus Expression System following manufacture’s instruction (Invitrogen, Carlsbad, CA, Cat#12562-054). The RBM20 concentration was determined by Bradford protein assay prior to enzymatic digestion. For each enzymatic digestion, approximately 50 μg of purified RBM20 was reduced in 6 mM DTT solution at 37 °C for 3 h and then alkylated in 5 mM iodoacetic acid at 37 °C for 30 min. The reduced and alkylated RBM20 was then digested with an enzyme to protein ratio of either 1:100 (for Lys-C, Glu-C, and Asp-N) or 1:200 (for trypsin) to produce large peptides with mass range of 3-20 kDa. The reactions were incubated in 25 mM ammonium bicarbonate buffer at pH 8 for 30 min at 37 °C and then quenched by acidifying the solution with 2 μL 98% formic acid (FA). 10 kDa molecular weight cut off (MWCO) filters and 0.1% FA in water were used to remove salt and small peptides from proteolytic products. The desalted large peptide mixture was separated by a home-packed PLRP column (PLRP-S, 200 mm length x 500 μm i.d., 10 μm particle size, 1,000 Å pore size, Agilent) in a 63-min gradient from 10% to 90% mobile phase B (mobile phase A: 0.1% FA in water; mobile phase B: 0.1% FA in 1:1 acetonitrile:ethanol) in a nanoACQUITY liquid chromatography (LC) system (Waters, Milford, MA, USA). The nanoACQUITY LC system was coupled with an impact II quadrupole-time-of-flight (q-TOF) mass spectrometer (Bruker, Bremen, Germany) for online mass spectrometry (MS) and tandem MS (MS/MS) analysis ^37-38^. Mass spectra were collected at 1 Hz over the range of 200-3000 *m/z*. The isolation window for online MS/MS of auto collision induced dissociation (CID) was set to 4 *m/z*. The collision DC bias was set from 18 to 50 V for CID with nitrogen as collision gas.

For offline high-resolution MS/MS analysis of phosphorylated peptides, the eluates from LC separation were collected every 1 minute from retention time of 15 min to 45 min. Then the peptide fractions were analyzed by a 12-T SolariX Fourier-transform ion cyclotron resonance (FTICR) mass spectrometer (Bruker, Bremen, Germany) equipped with electron capture dissociation (ECD)^39^ The samples were introduced into the mass spectrometer using an automated chip-based nano-electrospray ionization (nano-ESI) source (Triversa NanoMate, Advion Bioscience, Ithaca, NY, USA). The mass spectra were collected over the range of 200-3000 *m/z* with 2 M transient size (1.153s) and 28%-30% excitation power. The isolation window for ECD fragmentation was set to 2-3 *m/z* and the ECD bias was from 0.8 to 1.0 V. The MS/MS spectra was the sum of 200-500 transients.

The online LC-MS and MS/MS data were processed and analyzed by DataAnalysis software from Bruker Daltonics. Mass spectra of peptide profiling were deconvoluted using the Maximum Entropy algorithm in the DataAnalysis software. For online MS/MS data, the msalign file of each MS/MS spectrum was output from the DataAnalysis software for identification and characterization of RBM20 peptides using MS-Align+^40^. The cutoff of E-value and P-value was set to E-10 for the confident identification of peptides. For offline MS/MS data, the mass and charge lists of fragmentation ions were output from DataAnalysis software for peptide sequence identification using MS-Align+. In-house developed MASH Suite Pro^41-42^ was used for the manually validation of fragment ions and localization of phosphorylation sites. A minimum fit of 60% and S/N threshold of 3 were set for peak picking. Fragment ions including *c, c*-1, *z*^*•*^, *and z*^*•*+1^ ions were validated within 10 ppm mass error. The reported masses for the RBM20 peptides and the corresponding fragment ions are all monoisotopic masses.

### Confocal imaging experiments

Hela cells were transfected with pEGFP-C1-8his-RBM20 WT/mutations plasmids, and 48 hours later cells were fixed with methanol for 20 min at −20°C. After 5 times washing with 1× PBS, cells were mounted in DAPI-containing mounting medium (Vector Laboratories) and observed with a Zeiss LSM-710 confocal microscope with 405 nm and 488 nm laser lines.

### Western blot

Western blotting analysis was performed as described previously^35-36^. Briefly, Protein lysed from cells and tissues were separated by SDS-PAGE gel and transferred onto a PVDF membrane. The membrane was probed with antibodies against AKT2 (Santa Cruz, CA, Cat#sc-81436), CLK1 (Santa Cruz, CA, Cat#sc-515897), SRPK1 (Santa Cruz, CA, Cat#sc-100443), GAPDH (Cell Signaling, MA, Cat#14C10) was served as the protein loading control.

### Data analysis

GraphPad prism software was used for statistical analysis. Results were presented as mean ± SEM. Statistical significance for each variable was estimated by the unpaired t-test (two-tailed) or the one-way analysis of variance (ANOVA) followed by a Tukey’s post-hoc analysis. Significance was considered as probability values of P<0.05 indicated by one asterisk, P<0.01 indicated by two asterisks, and P<0.001 indicated by three asterisks.

## Results

### Phosphorylation identification of RBM20 with middle-down MS

SR domain in a splicing factor is frequently phosphorylated and its phosphorylation plays a key role in pre-mRNA splicing. RBM20 contains one SR domain, however, the phosphorylation status of RBM20 SR domain remains unknown. To identify the phosphorylation of RBM20, we purified RBM20 from wild type (WT) and knockout (KO) fresh rat heart tissues respectively through immunoprecipitation (IP) using anti-RBM20 antibody. Purified RBM20 was subject to western blot and pro-Q diamond staining, a dye indicating phosphorylation of a protein. A protein band with pro-Q diamond staining was observed in WT heart (*RBM20*^*+/+*^) but not in KO heart (*RBM2*^*-/-*^) (**Fig. 1A**). The size of the protein band is the same as RBM20 band identified with western blot against anti-RBM20 antibody (**Fig. 1B**). Interestingly, two RBM20 bands appear in WT heart with western blotting analysis, suggesting potential PTMs lead to partial RBM20 shifting (**Fig. 1B**). To further confirm what we observed with RBM20 expressed in fresh heat tissues, we expressed and purified RBM20 with the *in vitro* expression system in both prokaryotic (*E. coli*) and eukaryotic (sf9 insect) cells. Coomassie staining shows that RBM20 protein purified from both insect sf9 cells and *E. coli* (**Fig. 1C**), whereas pro-Q diamond only stains RBM20 protein expressed in insect cells, but not in *E. coli* (**Fig. 1D**). These results suggest that RBM20 is highly likely to be phosphorylated. To unambiguously validate the phosphorylation status of RBM20, middle-down MS was performed to identify the phosphorylated residues with purified RBM20 protein from insect sf9 cells. One of the representative results from MS analysis is shown in **Figure 1E** and other MS results are presented in supplementary figures (**Fig. S1-11**). Total sixteen phosphorylation sites were identified with four of them (S638, S640, S643, and S645) on SR domain and the rest of them located at S652, S729, S789, S879, S881, S999, S1034, S1046, S1057, S1096, S1190 and S1192 (**Fig. 1F and S12**). Eleven RBM20 mutations have been found in human patients with dilated cardiomyopathy (DCM), and five residues are substituted due to the genetic mutations in SR domain^10-16^ (**Fig. 1F**). Among them, two residues S635 (S638 in rat) and S637 (S640 in rat) are phosphorylated and substituted by Alanine (A) and Glycine (G) respectively in human patients with DCM, suggesting that mutations S to A or G may reduce the phosphorylation level on SR domain, and thus interrupt the splicing function of RBM20.

**Figure 1.**
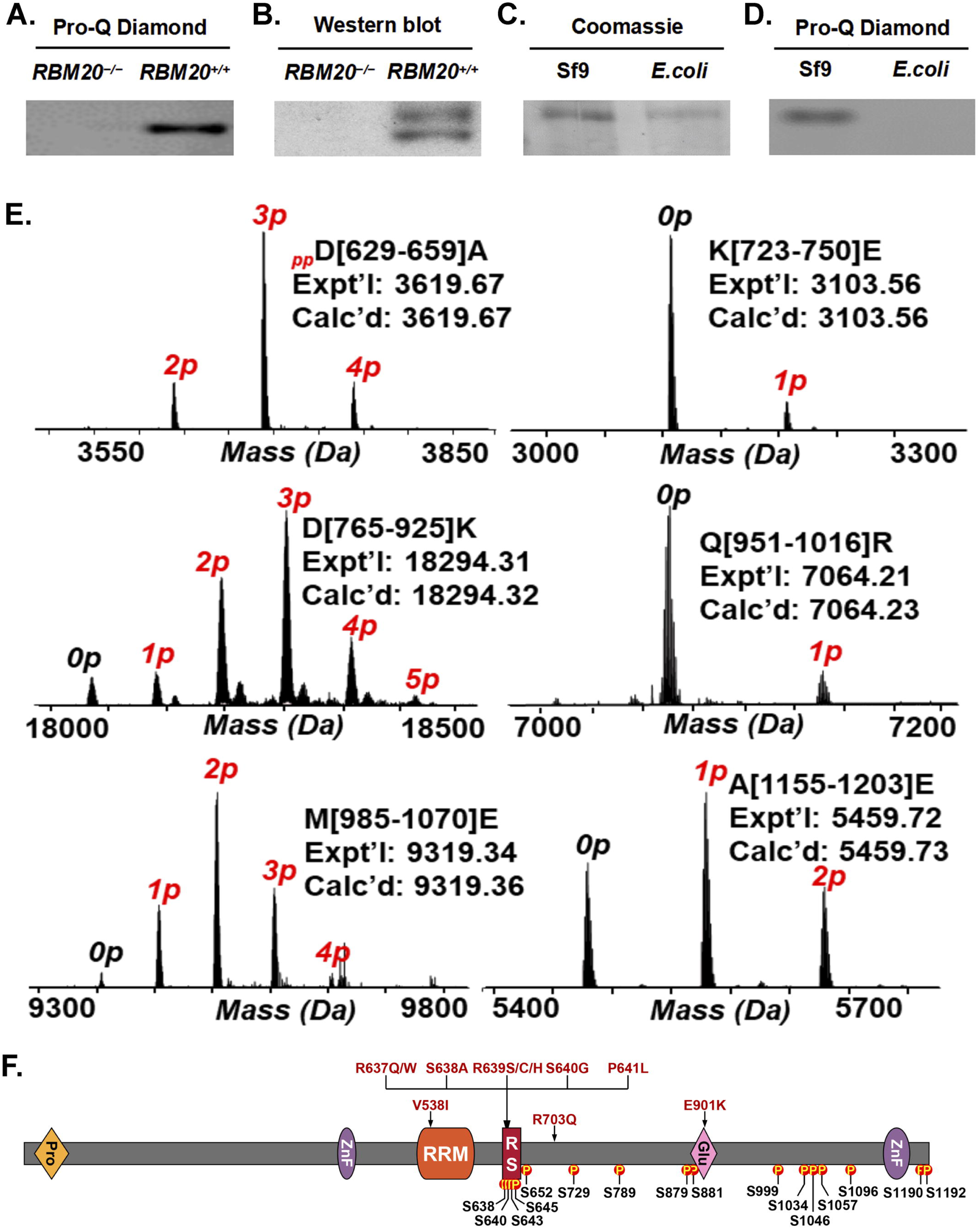
Characterization of RBM20 phosphorylation. **A**, ProQ diamond staining of RBM20 purified from WT and KO heart tissues; **B**, Western blot verification of RBM20 purified from WT and KO heart tissues; **C and D**, Coomassie and proQ diamond staining of RBM20 protein expressed in sf9 insect cells and *E. coli*; **E**, Mass spectra of six representative phosphorylated RBM20 peptides, D[629-659]A, K[723-750]E, D[765-925]K, Q[951-1016]R, M[985-1070]E, and A[1155-1203]E. Peptide sequences were assigned based on the MS/MS data from either CID or ECD fragmentation. The mass difference between each peak is 79.97 Da, indicating the occurrence of phosphorylation. The number on each peak shows the number of phosphorylation sites. Expt’l is the experimental monoisotopic mass based on data obtained from MS experiments. Calc’d represents the calculated monoisotopic mass based on amino acid sequences; **F**, Schematic diagram of phosphorylation sites on RBM20 and known mutations on RBM20 highlighted in red. *RBM20*^*-/-*^, knockout; *RBM20*^*+/+*^, wild type; Pro, proline; ZnF, Zinc finger domain; RRM, RNA recognition motif; RS, arginine/serine; Glu, Glutamic Acid. P, phosphorylation.

### Phosphorylation on SR domain is critical in the regulation of pre-mRNA splicing

RBM20 is a muscle specific splicing factor that plays a major role in titin pre-mRNA splicing^6^. To determine the impact of RBM20 phosphorylation on pre-mRNA splicing, sixteen phosphorylation sites were mutated to non-phosphorylatable A except for S640 that was mutated to G to mimic mutation in human patients with DCM. RBM20 mutations were constructed into pEGFP-C1 vector. These mutated constructs were co-transfected into HeLa cells with titin mini-gene plasmid including exons 64 to 70 (**Fig. 2A**). WT RBM20 was used as positive control and empty plasmid was used as negative control (NC). RNA was purified from each treatment 48h after co-transfection and RT-PCR were performed to detect splicing pattern of titin mini-gene. In the absence of RBM20 (NC), a single smaller band can be detected while in the presence of RBM20 (WT), two larger bands appear and total three bands can be detected (**Fig. 2C-F**). Fourteen out of sixteen mutated RBM20 show the same splicing pattern as WT RBM20 (**Fig. 2C-E**), however, two mutations S638A and S640G indicate the same splicing pattern as the negative control (**Fig. 2F**). These two mutations are located in the SR domain. They have also been identified in human patients with DCM (**Fig. 1F**). This *in vivo* cell splicing assay revealed that mutations on SR domain, particularly, on phosphorylation sites eliminate the splicing function of RBM20. To further confirm the observation, a dual luciferase splicing reporter assay was performed with individual RBM20 mutations. The reporter construct contains exons 4 to 13 in titin PEVK region as described previously^6^. High ratios of firefly to renilla luciferase means suppression of splicing and vice versa (**Fig. 2B**). The exact same results were observed with the RT-PCR results. Only two mutations S638A and S640G suppress the splicing function of RBM20 (**Fig. 2G**), which proves the concept that phosphorylation on SR domain plays a key role in pre-mRNA splicing.

**Figure 2.**
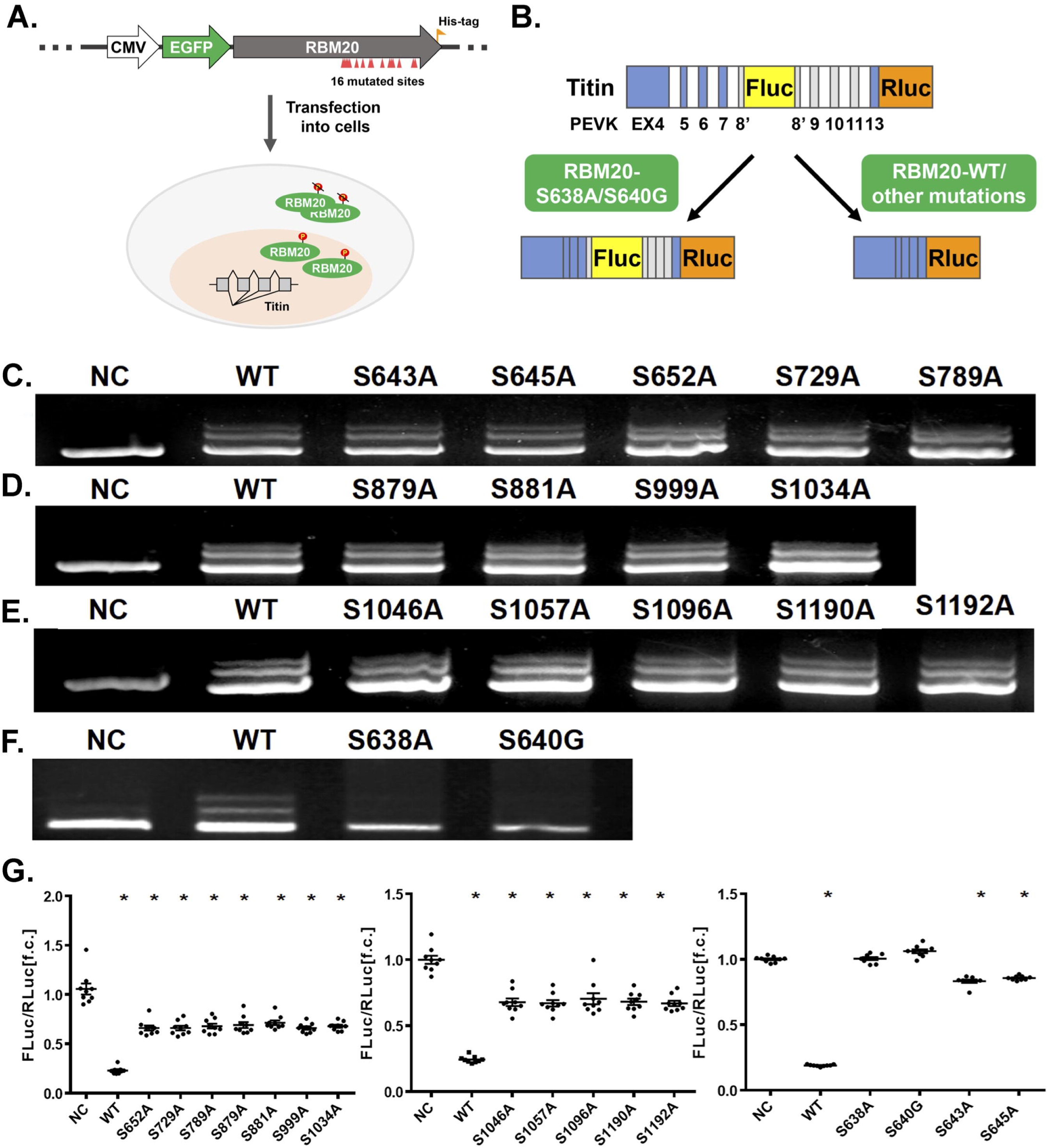
Impact of mutations on RBM20 phosphorylation sites on TTN splicing. **A**, Schematic diagram of RBM20 constructs and co-transfection in HeLa cells. **B**, Schematic diagram of TTN mini gene (PEVK Ex4-13) construct with dual luciferases and principle of detection; **C-F**, PCR detection of TTN mini gene splicing pattern; **G**, Dual luciferase splicing assay with PEVK Ex4-13 splice reporter in HEK293 cells. Ex, exon; WT, RBM20 wild type; Fluc, firefly luciferase; Rluc, renilla luciferase; NC, negative control with empty plasmid; Mean±SEM (n=3); * P<0.05.

### Mutations S638A and S640G on SR domain facilitate RBM20 trafficking from the nucleus to the cytoplasm and protein aggregation in the cytoplasm

RBM20 is a nuclear protein. Under stress conditions, the proteins can be relocated from the nucleus to the cytoplasm and form stress granules^43-46^. To validate if this is the case for mutations in SR domain, especially for mutations on phosphorylation sites, we transfected 16 mutations as well as a double mutation of S638A/S640G into HeLa cells. Confocal fluorescent microscope was applied to detect the localization of RBM20. The results revealed that only mutations S638A and S640G as well as double mutations S638A/S640G facilitate RBM20 trafficking from the nucleus to the cytoplasm (**Fig. 3A**). The rest of the fourteen mutations retains the RBM20 in the nucleus as the same as WT RBM20 (**Fig. 3A**). Most interestingly, we observed the granules formation or protein aggregation in the cytoplasm with mutations S638A and S640G as well as the double mutations S638A/S640G (**Fig. 3A**). These results suggest that only mutations on phosphorylation sites in SR domain facilitate RBM20 trafficking and protein aggregation, which may imply new role of RBM20 in the progression of cardiac dysfunction.

**Figure 3.**
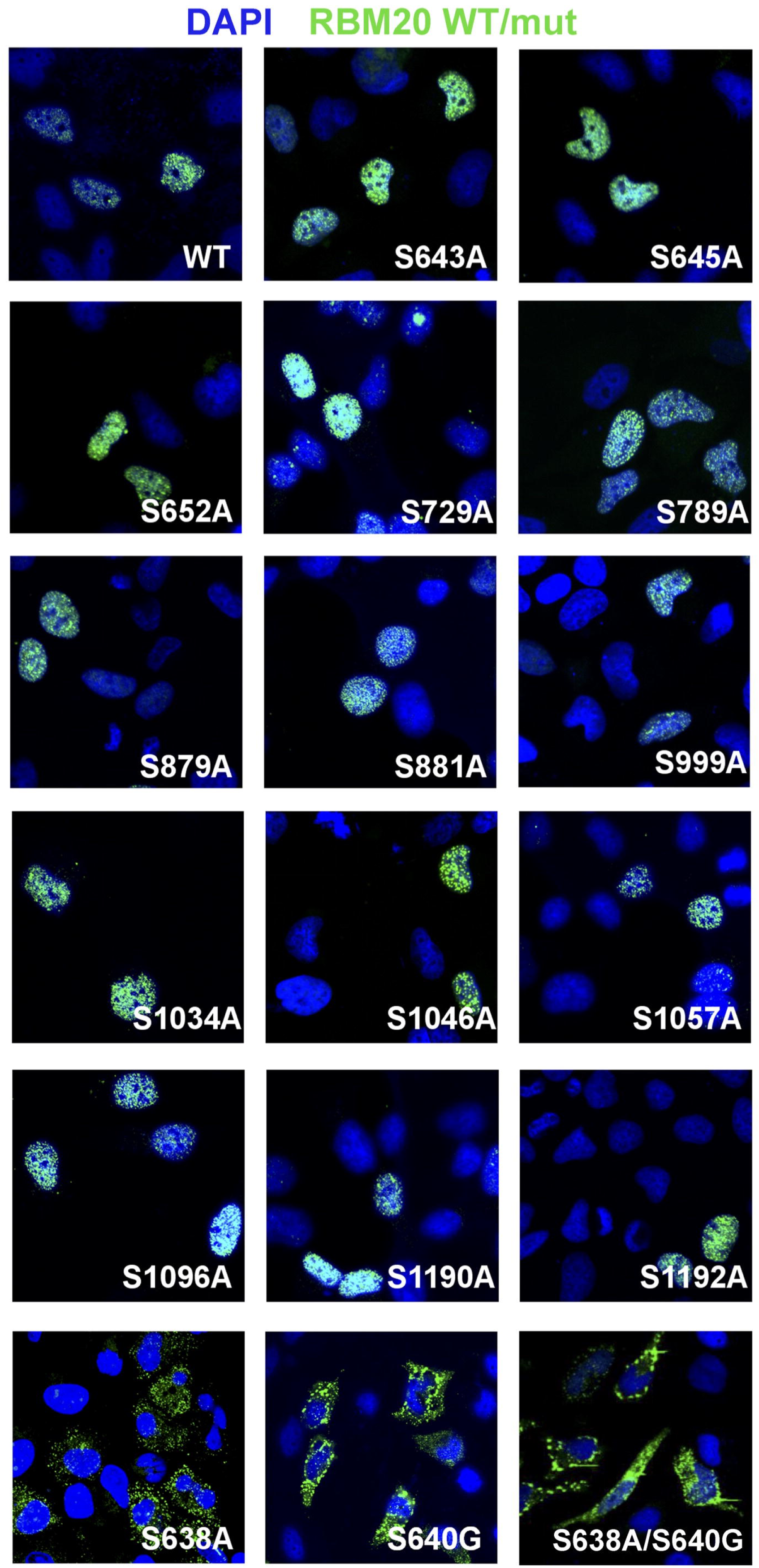
Confocal imaging of RBM20 mutations in HeLa cells. HeLa cells were transfected with RBM20 or RBM20 mutations in GFP fusion plasmid respectively. Localization of RBM20 was visualized by confocal microscope. Nucleus was stained with DAPI. WT, wild type; mut, mutation.

### AKT, CLK and SRPK are Kinases phosphorylating RBM20

AKT, CLK and SRPK kinase families have been shown to commonly phosphorylate the SR domain in splicing factors^24-26^. To test whether these kinase families phosphorylate RBM20, AKT2, CLK1 and SRPK1 kinases plasmids were co-transfected with RBM20 mutation constructs respectively in HeLa cells. WT RBM20 was used as control. Cells were harvested and lysate 48 h post co-transfection. Proteins from each treatment were subject to western blotting using antibody against anti-phospho-RBM20 (Gift from Dr. Kuroyanagi Hidehito)^47^ and anti-pan-RBM20. The results reveal that phosphorylation level of WT RBM20 is significantly increased by all three kinases when compared to WT RBM20 without kinases treatment (**Fig. 4A-C**). In cell lysates co-transfected with RBM20 mutations, S638A or S640G or S638A/S640G, phosphorylation level of RBM20 is significantly reduced when compared to WT RBM20, meaning that these two mutations are the major contributors to RBM20 phosphorylation. Furthermore, phosphorylation level of RBM20 with S638A has no difference between treated and untreated cell lysates, suggesting S638 could be the major residue phosphorylated by kinases AKT, CLK and SRPK. Phosphorylation level of RBM20 with S640G mutation is significantly decreased without kinase treatment, however, it can be restored by kinase treatment (**Fig. 4A-C**). Interestingly, double mutations S638A and S640G almost fully eliminate phosphorylation of RBM20 with or without kinase treatment, further confirming that S638 and S640 are important serine residues for RBM20 phosphorylation by these three kinases. S638 is a primary target of kinases AKT, CLK and SRPK, while S640 is perhaps primarily phosphorylated by other unknown kinases. Nevertheless, AKT2 seems having the lowest phosphorylation capacity on S638. CLK and SRPK have stronger phosphorylation possibility on S638 residue, but CLK seems also phosphorylating phosphor-sites of RBM20 other than S638 and S640. The *in vitro* kinase assay further confirmed that all three kinases phosphorylate RBM20 with lower phosphorylation level by AKT (**Fig. 4D**), which is consistent with the *in vivo* kinases assay. Phosphorylation level has also been validated by LC-MS and LC-MS/MS analysis of RBM20 purified from the *in vitro* kinases assay with these kinases (**Fig. 4E and Fig. S13**). The results indicate that RBM20 treated with CLK1 and SRPK1 kinases has higher phosphorylation level than that treated without kinases and further confirmed that Akt is not a primary kinase for RBM20 phosphorylation (**Fig. 4E**).

**Figure 4.**
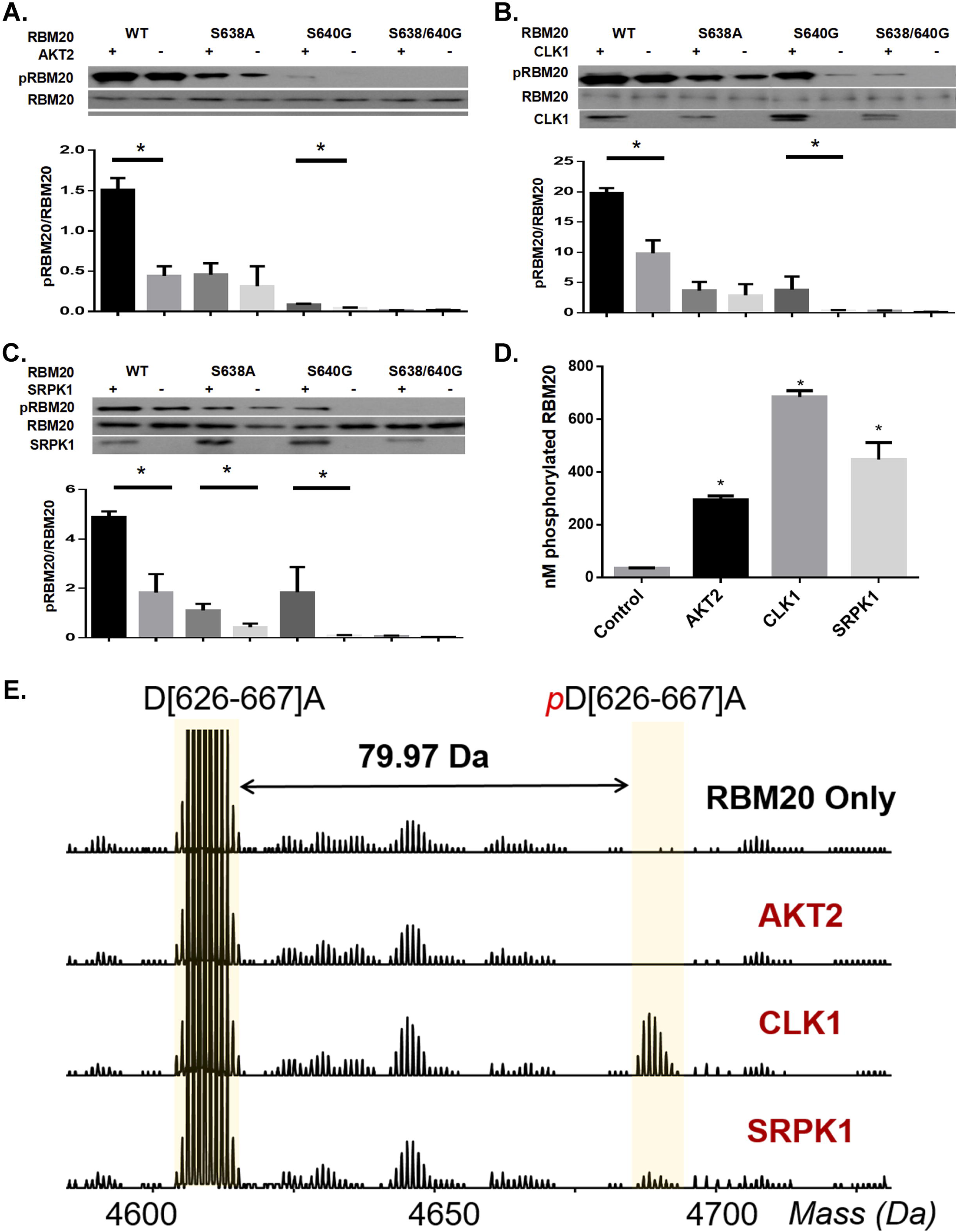
The *in vivo* and *in vitro* protein kinase assays. **A-C**, Western blotting with anti-phospho-RBM20 and anti-pan-RBM20 using protein preparations from HeLa cells with transfection of RBM20 mutations or co-transfection of RBM20 mutations and kinases (AKT2, CLK1 and SRPK1) respectively. WT RBM20 was used as control. **D**, *in vitro* kinase assay; **E**, LC-MS analysis with samples from the *in vitro* kinase assay; mean ± SEM (n=3); * P< 0.05.

### Kinases AKT2, CLK1 and SRPK1 interact with RBM20 and regulate mRNA splicing in HeLa cells

RBM20 phosphorylation by kinases AKT, CLK and SRPK suggests potential interaction between RBM20 and kinases. To determine the protein-protein interaction, RBM20 plasmid with his-tag was co-transfected with individual kinase constructs respectively in HeLa cells. Cells were harvested 48 h after co-transfection and the protein lysates from the cells were subject to co-immunoprecipitation (co-IP). Anti-his tag antibody was used to capture RBM20 protein complex that is subsequently subject to western blotting with anti-RBM20, anti-SRPK1, anti-CLK1, and anti-HA-AKT2 separately. Western blotting analysis showed that SRPK1 (**Fig. 5A**), CLK1 (**Fig. 5B**), and AKT2 (**Fig. 5C**) are captured by RBM20. On the other hand, anti-SRPK1, - CLK1 and –HA-AKT2 were used as the bait to capture the prey RBM20. The results indicated that all three kinases could also pull down RBM20 (**Fig. 5D-F**). Controls without the antibody conjugated on beads did not capture any target proteins (**Fig. 5A-F**). To further validate if these three kinases can regulate titin pre-mRNA splicing through changes of RBM20 phosphorylation level, RBM20 WT and mutations were co-transfected with titin mini-gene and individual kinase constructs respectively in HeLa cells. Cells were harvested 48 h after co-transfection and total RNA was purified from these cells. RT-PCR was performed to detect the changes of splicing pattern of titin mini-gene. Three bands can be detected with WT RBM20 in the absence of kinases (NC), while only one larger band was detected in the presence of each individual kinase (**Fig. 5G**). A single smaller band was detected with S638A mutated RBM20 in the absence of kinases, whereas it was switched to two larger bands in the presence of CLK1 and SRPK1 and to the largest band in the presence of AKT2. The similar splicing pattern was observed with S640G and double S638A/S640G mutated RBM20 respectively. The only difference is that the single smallest band was completely switched to a single largest band in the presence of kinases CLK1 and SRPK1 by comparing to mutation S638A (**Fig. 5G**). These results suggest that CLK1, SRPK1 and AKT2 phosphorylate RBM20 through interaction with RBM20 and regulate titin pre-mRNA splicing via changing RBM20 phosphorylation level.

**Figure 5.**
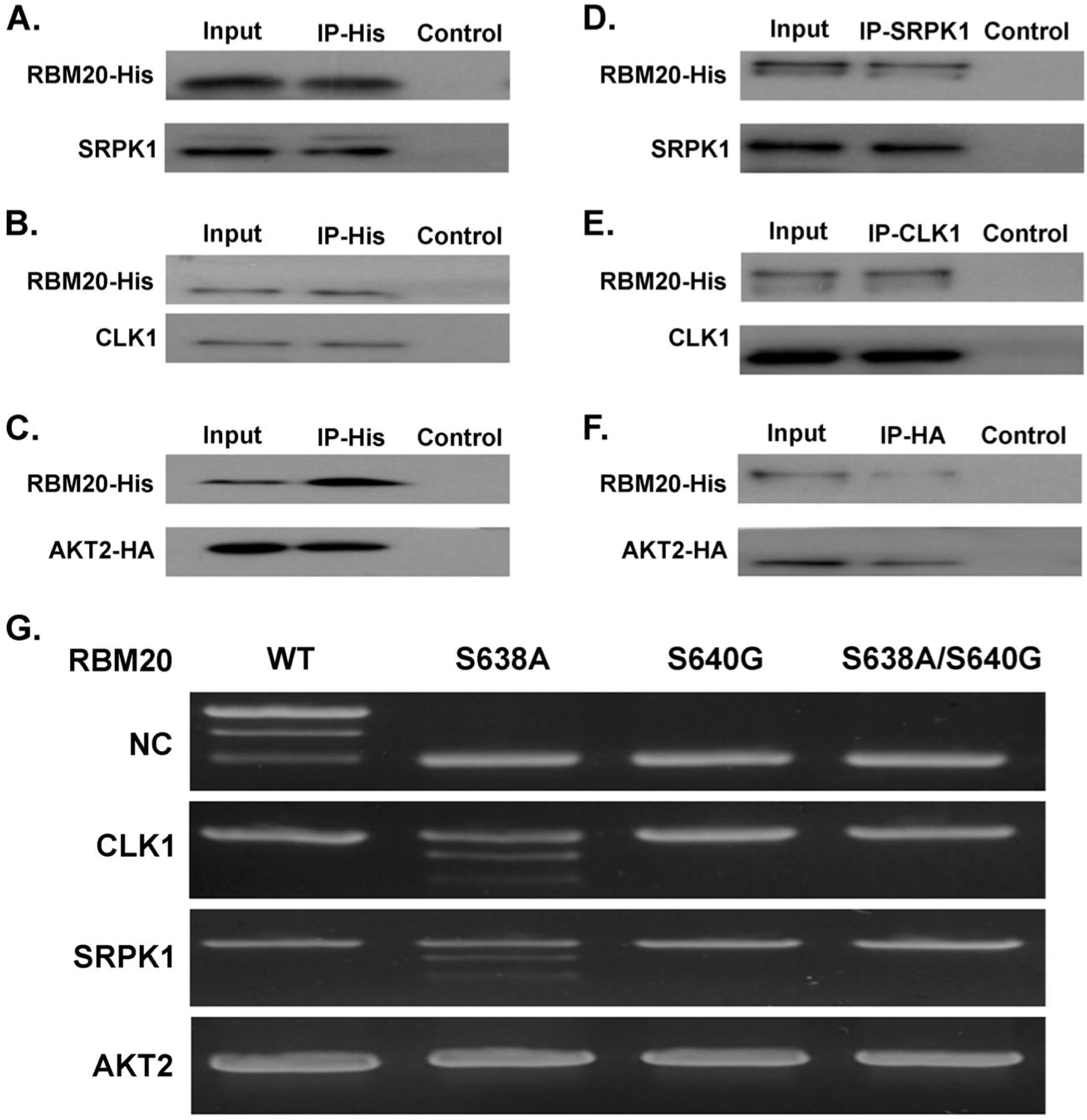
Co-immunoprecipitation (co-IP) and *in vivo* cell splicing assay. **A-C**, Western blotting with protein samples from co-IP using RBM20-His antibody as bait; **D-F**, Western blotting with protein samples from co-IP using anti-SRPK1, anti-CLK1 and anti-AKT-HA as bait respectively; **G**, *in vivo* cell splicing assay with co-transfection of individual kinases, RBM20 WT and mutations and TTN minigene (Mid-Ig Ex 64-70) in HeLa cells. Input, total protein before co-IP; IP-His, elution from co-IP with anti-His antibody; Control, elution from co-IP without antibodies; NC, control without kinase transfection; WT, RBM20 wild type.

### Inhibition of Kinases CLK and SRPK reduces RBM20 phosphorylation and interrupts pre-mRNA splicing in NRCMs

Inhibitors SRPIN340 and TG003 inhibit activity of kinase SRPK1 and CLK1 respectively. To verify whether inhibition of endogenous kinases SRPK1 and CLK1 in primary cell cultures impacts on RBM20 phosphorylation and pre-mRNA splicing of its targets, primary NRCMs were cultured and treated with these two inhibitors. NRCMs were harvested at different time points (0 min, 5 min, 1h, 6h, 12h, 24h and 48h). Proteins from NRCMs at individual time points were subject to western blotting analysis using anti-phospho-RBM20 and anti-pan-RBM20. Phosphorylation level of RBM20 is decreased at about 5 min with further reduction through longer treatment (**Fig. 6A, 6B and 6D**). The vehicle control with DMSO treatment does not change the phosphorylation of RBM20 with time courses (**Fig. 6C and 6D**). Both titin and Camk2d pre-mRNAs are the substrate of RBM20. Inhibition of kinases SRPK1 and CLK1 increases the larger variants’ expression of both titin (t-1) and Camk2d (d-1) gene when compared to the control DMSO treatment (**Fig. 6E**).

**Figure 6.**
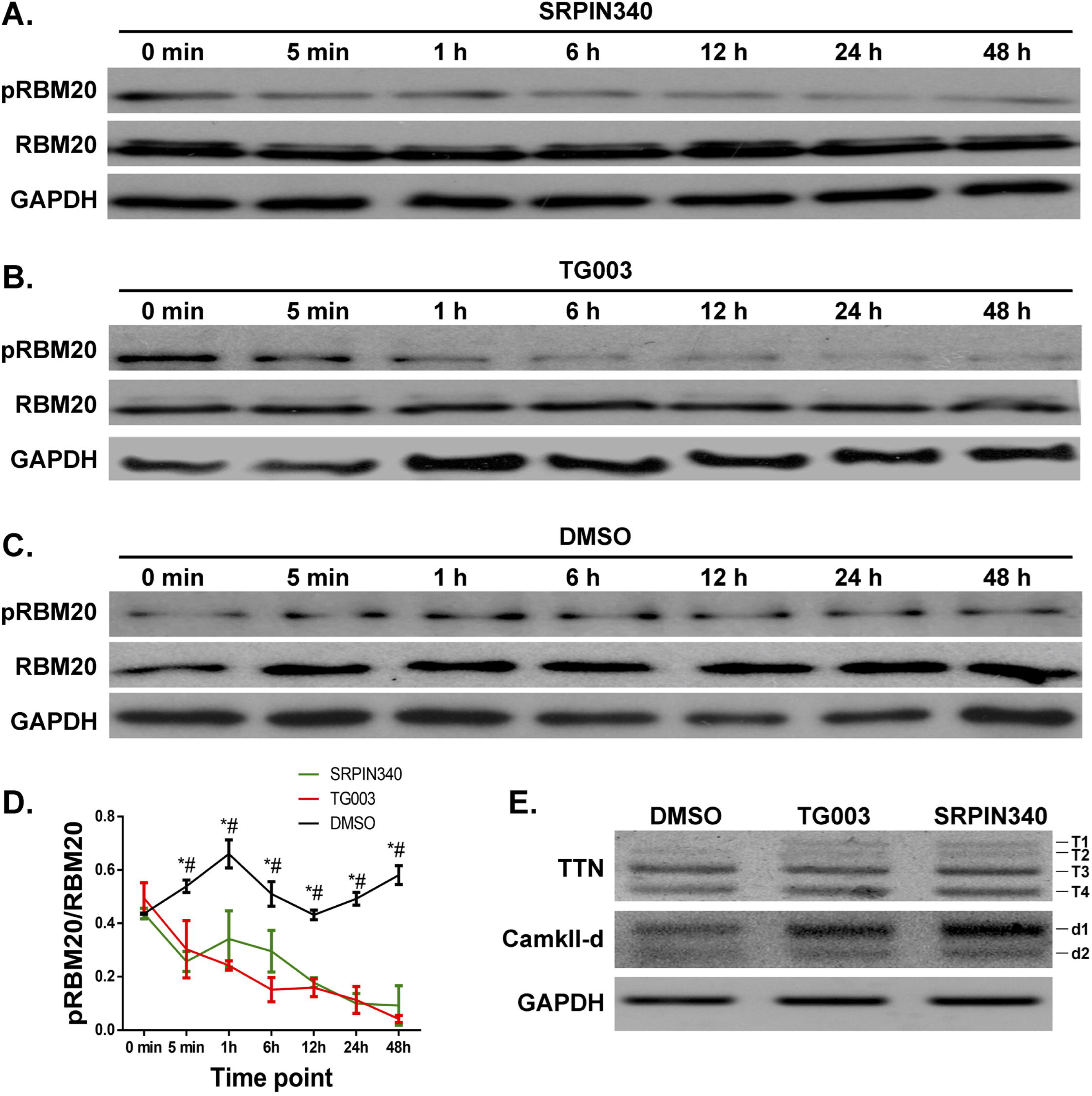
Impact of inhibition of SRPK1 and CLK1 on RBM20 phosphorylation and mRNA splicing. **A-C**, Western blotting against anti-phospho-RBM20 and pan-RBM20; Protein samples from NRCMs treated with inhibitors SRPIN340 or TG003 at different time points; vehicle DMSO was used as control. **D**, Quantification of western blotting results from A-C; **E**, Changes of splicing variants of TTN and CamkIId detected by RT-PCR. TTN variants, T1-T4; CamkIId variants, d1-d2. GAPDH, house keeping gene for RT-PCR and protein loading control for western blotting. Mean ± SEM (n=3); ^*/#^P< 0.05.

### AKT kinase regulates pre-mRNA splicing of RBM20 targets *in vivo*

Heart tissues from Akt2 KO and overexpression mice (**Fig. 7A and 7B**) were used to determine the phosphorylation of RBM20 and pre-mRNA splicing of RBM20 targets *in vivo*. Heart tissues collected from WT mice were used as control. Proteins prepared from these heart tissues were subject to western blotting analysis. RBM20 phosphorylation level is significantly reduced in Akt2 KO, whereas it is dramatically increased in Akt2 overexpression mice by comparing to WT (**Fig. 7A and 7C**). Interestingly, we observed that RBM20 expression is also affected by Akt level. Akt2 KO reduces RBM20 expression, whereas Akt2 overexpression increases RBM20 level (**Fig. 7A and 7D**). AKT regulating RBM20 expression has been confirmed in our previous publication^35-36^. Here, we report that AKT can also change the phosphorylation level of RBM20, and thus, downstream gene posttranscriptional process as shown in **Figure 7E**. By comparing to WT, Akt KO increases larger isoform expression d-1 and d-2, but no change was observed in titin splicing pattern. However, Akt overexpression promotes titin larger isoform expression (L), but not CamkIId (**Fig. 7E**). These results suggest Akt kinase regulates pre-mRNA splicing of RBM20 targets through potential different mechanisms which need be further studied.

**Figure 7.**
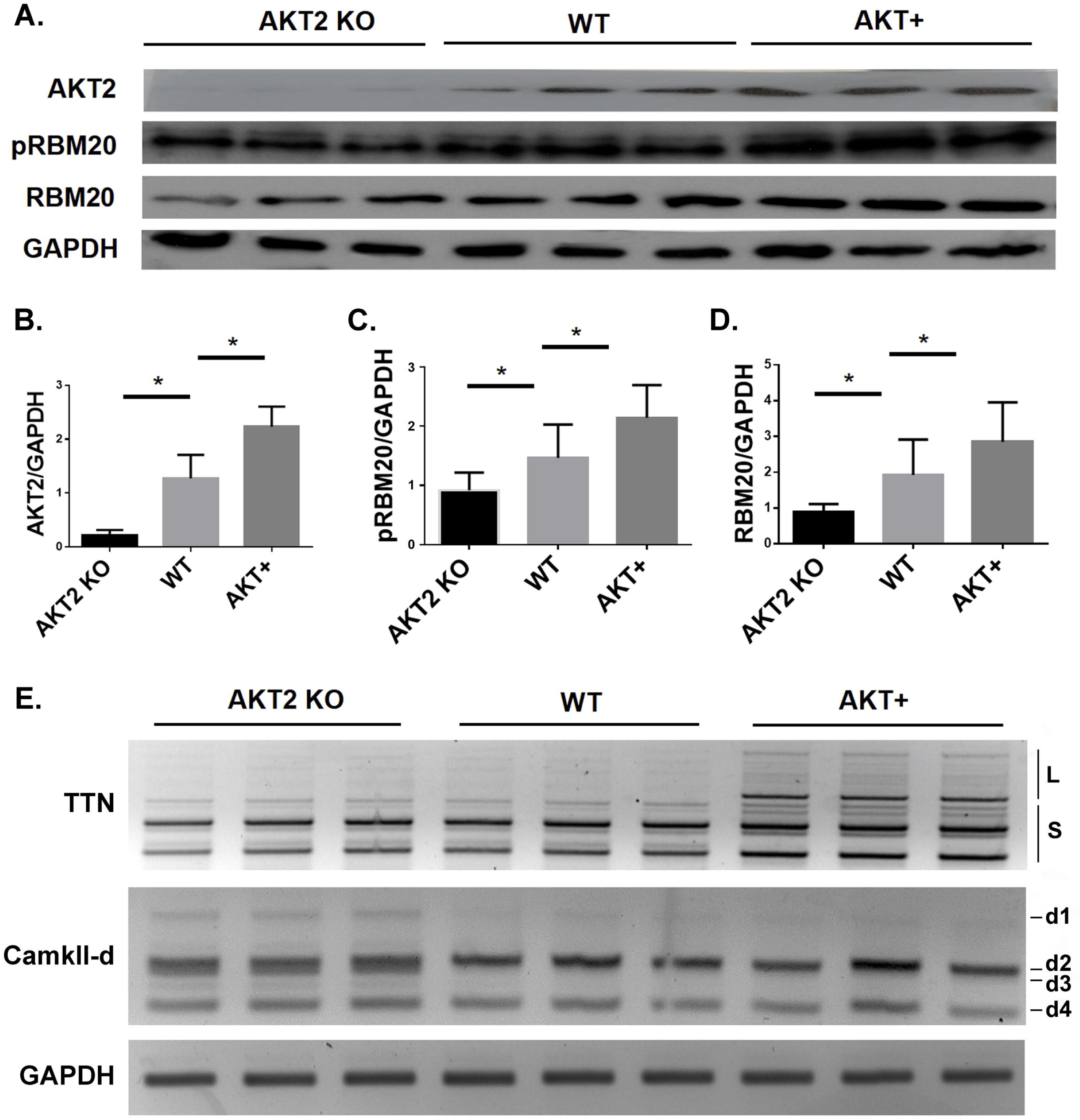
RBM20 phosphorylation and splicing alteration in AKT2 KO and overexpression mice. **A**, Western blot analysis of RBM20 phosphorylation, RBM20 expression and AKT2 expression in AKT2 KO and overexpression mice respectively. WT mice were used as control; **B**, Quantification of AKT2 expression level; **C**, Quantification of RBM20 phosphorylation level; **D**, Quantification of RBM20 expression level; **E**, Detection of TTN and CamkIId splicing alteration using RT-PCR. KO, knockout; WT, wild type; AKT+, AKT overexpression; L, a class of larger TTN variants; S, a class of smaller TTN variants; d1-d4, camkIId variants; GAPDH, house keeping gene for RT-PCR and protein loading control for western blotting; mean ± SEM (n=3); * P< 0.05.

## Discussion

Our salient findings revealed that RBM20 protein can be phosphorylated on sixteen serine amino acid residues. Four out of sixteen phosphorylated serine residues are located in SR domain. Individual mutations of fourteen serine to the unphosphorylatable A do not affect splicing function of RBM20. However, the rest two mutations S638A and S640G on SR domain alter the splicing pattern of RBM20 substrates such as titin and camk2d. Most intriguingly, mutations S638A and S640G facilitate RBM20 trafficking from the nucleus to the cytoplasm and form protein aggregation, while mutations on other fourteen phosphorylation sites retain RBM20 in the nucleus. Further, we found that phosphorylation sites S638 (S635 in human) and S640 (S637 in human) can be phosphorylated by kinases AKT, CLK, and SRPK. S638 is highly phosphorylated by CLK and SRPK, but slightly phosphorylated by AKT, while S640 is perhaps primarily phosphorylated by other kinases. *In vitro* studies with inhibition of CLK and SRPK and *in vivo* studies with AKT KO and overexpression mice validated that kinases AKT, CLK, and SRPK regulate RBM20 phosphorylation level, and thus, pre-mRNA splicing.

Pre-mRNA splicing is a key mechanism for protein diversity as well as gene regulation that involves dynamic interactions of protein-RNA and protein-protein complexes^48^. In this tightly regulated process, SR proteins play a critical role in modulating the binding and assembly of the spliceosome complex and thereby determining the splicing site selection^48^. SR proteins are a group of RNA binding proteins containing an enriched dipeptide SR repeats, also defined as SR domain. SR domain can mediate protein-protein and protein-RNA interaction during spliceosome assembly ^48^. Splicing factors are a subfamily of SR proteins and act as the *trans* effects in the regulation of pre-mRNA splicing. Mutations disrupting the pre-mRNA splicing have been found deleterious to human diseases^49^. A wide range of disorders as well as heart disease are caused by mutations affecting splice consensus sites (*cis* effects) ^50^. However, mutations in *trans*-acting factors such as splicing factors have not been broadly studied, particularly not in heart diseases^51^. Human genetic studies have shown that mutations in SR domain of RBM20 protein are associated with DCM, young age at diagnosis, end-stage heart failure, and high mortality^7-13^. Two of these mutations are S635A (S638A in rat) and S637G (S640G in rat) that are identified as phosphorylation residues in this study. Previous studies indicate that phosphorylation and dephosphorylation of SR domain is critical for numerous steps in subcellular localization and splicing control of splicing factors^27-31^. RBM20 mutations from a phosphorylation residue (S) to an unphosphorylatable residue (A or G) on SR domain disrupt pre-mRNA splicing in cells and tissues. This study provides another evidence that dephosphorylation of a splicing factor through mutations on SR domain interrupts splicing control in a *trans* effect manner. Cardiac dysfunction caused by mutations in *trans*-acting factor RBM20 could be associated with aberrant splicing such as titin and camk2d, and thus pathophysiological changes.

On the other hand, many proteins such as SR proteins in the spliceosome complex continuously shuttle between the nucleus and the cytoplasm. Localization of these proteins can shift in response to stress signals as well as mutations^51-53^. Changes of distribution of splicing factors in the nucleus and in the cytoplasm could impact on splicing pattern as well as gene expression that are associated with a diverse set of diseases^49, 54^. SR protein trafficking/shuttling has been extensively studied in neurodegenerative diseases. For example, TDP-43 is one of the RNA binding proteins that regulates splicing of the pre-mRNA of *CFTR* (the gene mutated in cystic fibrosis). TDP-43 is normally located in the nucleus. Mutations in the TDP-43 gene (*TARDBP*) can facilitate cytoplasmic inclusion. Depletion of TDP-43 from the nucleus and protein accumulation in the cytoplasm are associated with both sporadic and familial forms of Alzheimer’s disease and amyotrophic lateral sclerosis (ALS), demonstrating a role for TDP-43 mutation in the pathogenesis of neurological diseases^55-58^. However, the pathogenic mechanism caused by the mutations is still unclear. It could be relevant to loss of splicing factor’s nuclear function or the toxicity of protein aggregation in the cytoplasm or the combination of both. Original study thought that RBM20 associated cardiac dysfunction is primarily resulting from the aberrant titin splicing^6, 14, 16^. Ours and recent studies suggest that RBM20 shuttling and protein aggregation in the cytoplasm could also be a major contributor to mutation caused cardiac dysfunction^47, 59^. Our current study provides a potential molecular and cellular mechanism that phosphorylation could play a key role in the pathogenesis of cardiac dysfunction.

Lastly, we previously reported that the change in stoichiometry of RBM20 is important to modulate pre-mRNA splicing patterns^6, 14^. Furthermore, we revealed that expression level of RBM20 can be regulated by the PI3K-AKT-mTOR kinase axis. Activation of the PI3K/AKT/mTOR signaling pathway via the hormone (insulin and thyroid hormone) stimulus activates p70S6K1 and inhibits 4E-BP1, and thus increases RBM20 expression and alters titin splicing patterns^35, 36^. Subsequent study observed that MAPK/ELK1 signaling pathway can also enhance transcription and expression of RBM20^60^. Notably, stoichiometric changes of RBM20 expression through the activation of signaling pathways slightly impact on titin gene splicing. This suggests that phosphorylation of RBM20 could play a predominant role in pre-mRNA splicing. Therefore, studying kinase signaling pathways could be the key in the development of novel treatment for the pathogenic progress of heart tissue induced by mutations in the splicing factors. Taken together, our present study revealed that RBM20 can be phosphorylated and for the first time identified the phosphorylation residues on RBM20. Phosphorylation and de-phosphorylation of RBM20 SR domain result in titin splicing transition. Also, mutations on phosphorylation sites leads to RBM20 shuttling from the nucleus to the cytoplasm and protein aggregation which could be a new role of RBM20 in the development of heart failure (**Fig. 8**). In summary, findings from our current research reveal the new mechanisms of mutation-caused pathophysiological changes in the heart.

**Figure 8.**
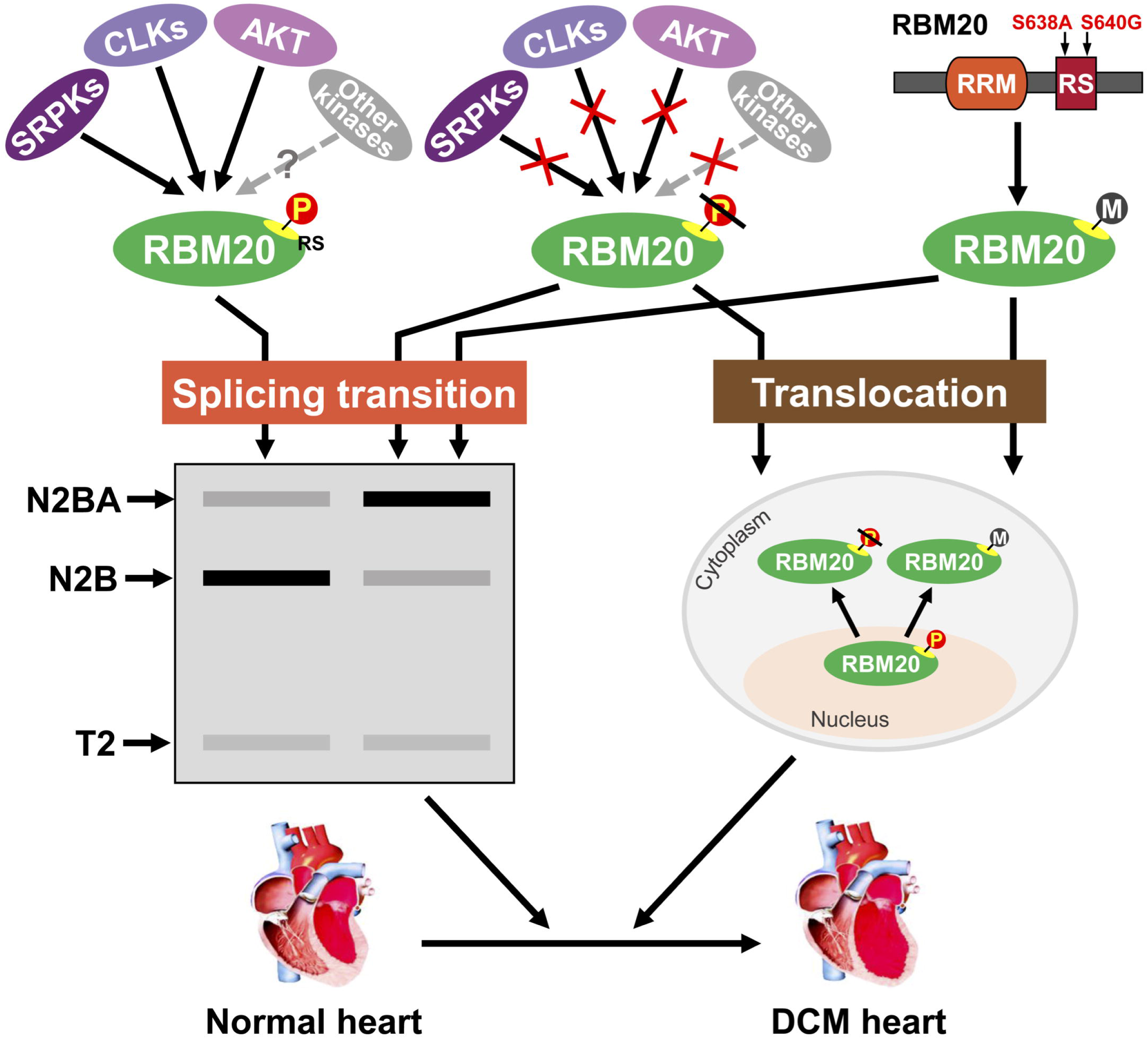
Schematic diagram of mechanisms of RBM20 phosphorylation on SR domain in the development of heart failure. Dephosphorylation and/or mutations of RBM20 SR domain facilitate titin splicing transition as well as protein shuttling, and thus result in cardiac dysfunction. Solid lines, identified kinases; dotted line, unknown kinases; P, phosphorylation; M, mutation. N2BA/N2B, titin isoforms; T2, titin degradation band

## Acknowledgement

We would like to thank to Dr. Kuroyanagi Hidehito at Tokyo Medical and Dental University and Dr. William Sellers for the generous gifts. This work was supported by the National Institute of General Medical Sciences of the National Institute of Health under award number P20GM103432; the Beginning Grand-in-Aid [16BGIA27790136 to WG] and the Transformational Project Award [19TPA3480072 to WG] from the American Heart Association; NIH R01 HL148733 to WG. YG would like to acknowledge NIH R01 grants, HL109810 and HL096971 and the high-end instrument grant S10OD018475.

---

**Figure S1.**
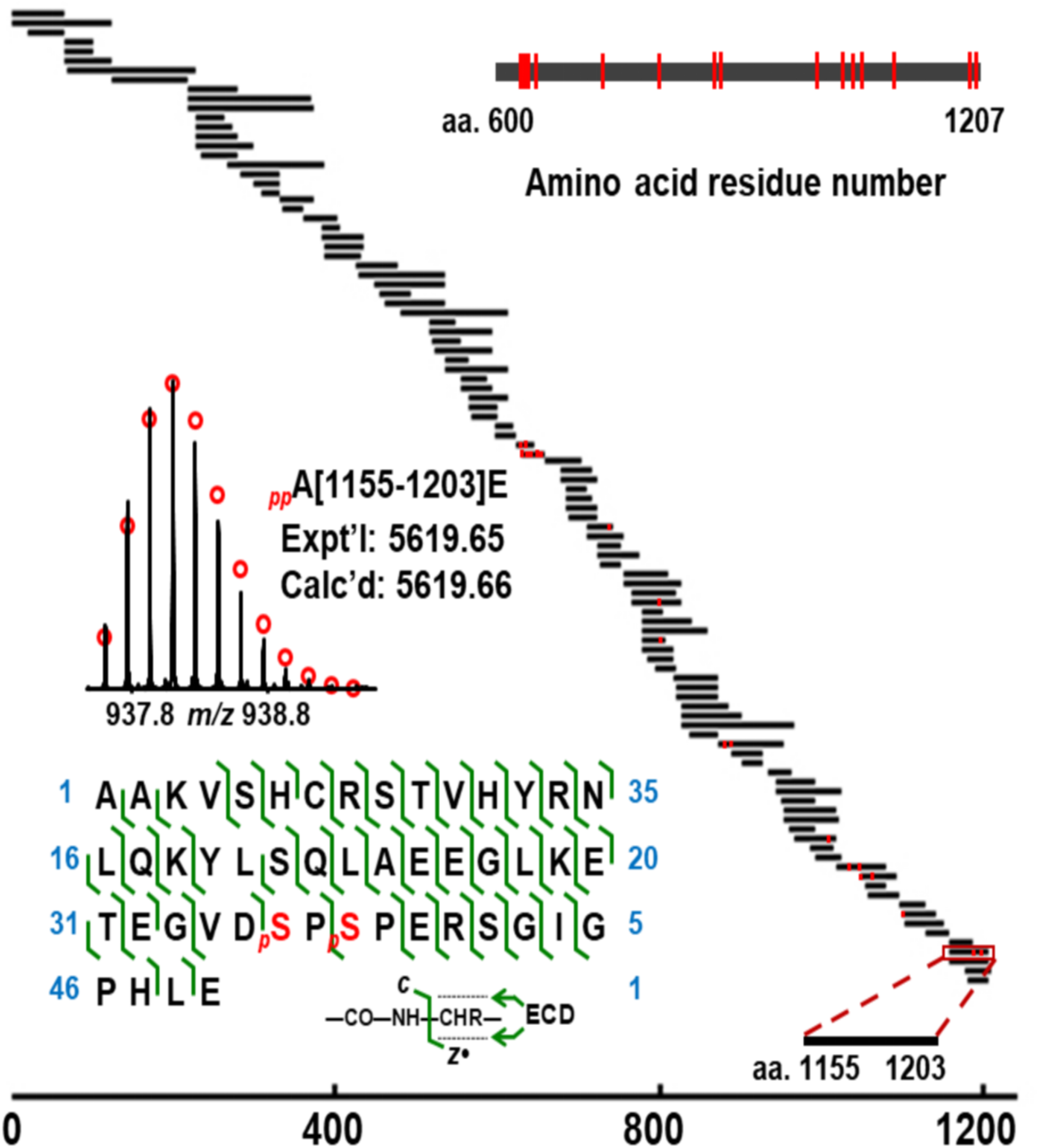
Schematic showing 100% sequence coverage of rat RBM20 and all phosphorylation sites characterized from the combination of limited Lys-C, Glu-C, Asp-N and trypsin digestions. Peptide sequence mapping was based on the MS/MS data from either CID or ECD fragmentation. The phosphorylation sites on peptides were marked in red. Inset: Left, a representative phosphopeptide, A[1155-1203]E, with confirmed sequence and phosphorylation sites Ser1190 and Ser1192 by ECD fragmentation. Right, all phosphorylation sites characterized on amino acid residues from amino acid residue 600 to 1207.

**Figure S2.**
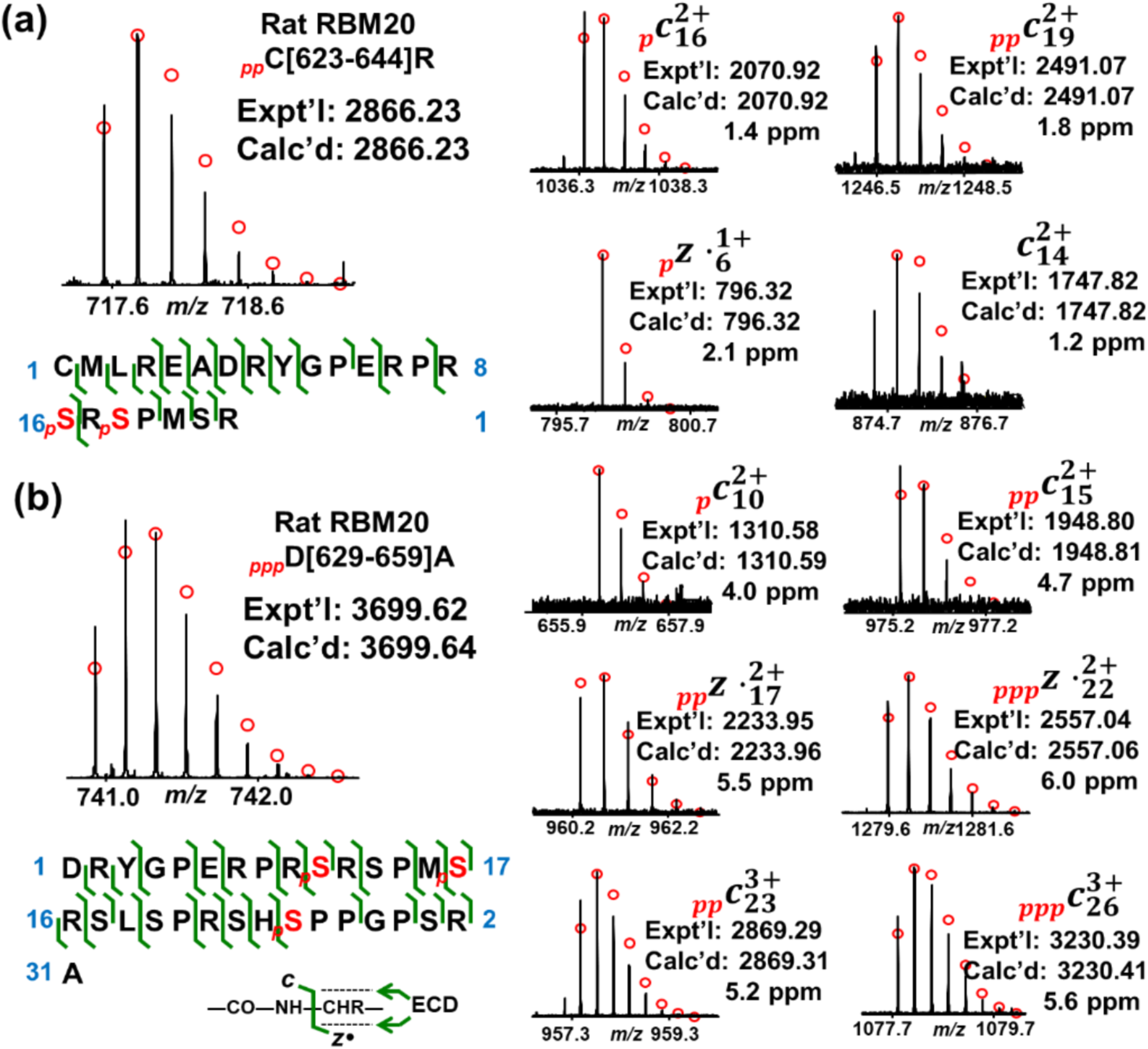
Identification of two phosphorylated peptides **(a)** C[623-644]R and **(b)** D[629-659]A with four phosphorylation sites, Ser 638, Ser 640, Ser 643, and Ser 652. Left, mass spectra of the precursor ion and sequence map of the phosphorylated peptide showing bond cleavages and phosphorylation sites. Cysteine carboxymethylation is considered in sequence mapping. The phosphorylation sites are highlighted in red. Right, representative fragment ions from ECD fragmentation. Fragment ions confirmed phosphorylation sites, Ser638 and Ser640, on peptide C[623-644]R and Ser638, Ser643, and Ser652, on peptide D[629-659]A.

**Figure S3.**
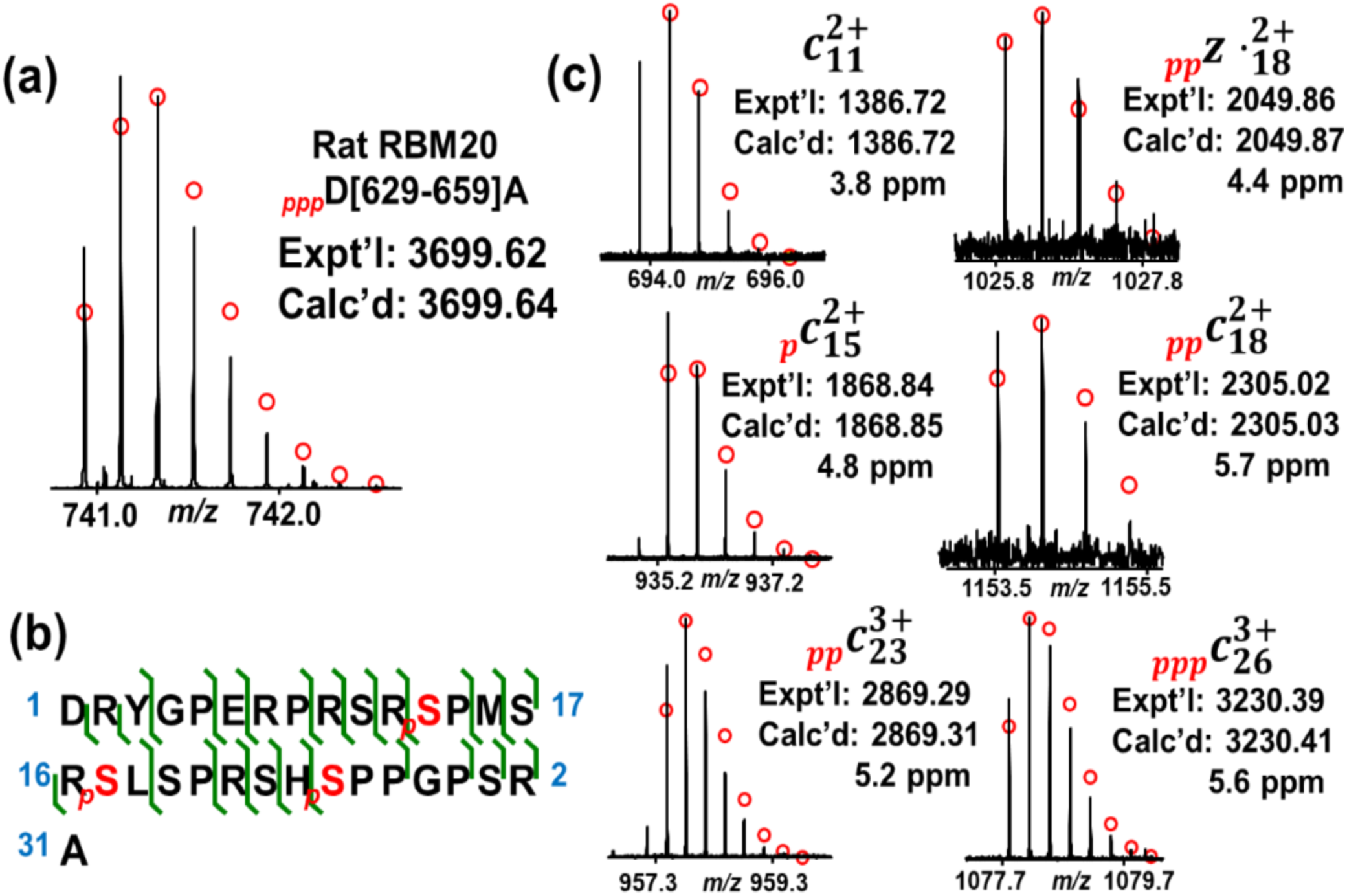
Identification of a phosphorylated peptide D[629-659]A with three phosphorylation sites, Ser640, Ser645 and Ser652, from Asp-N digestion of rat RBM20 (RBM20_Rat, UniProtKB-E9PT37). **(a)** Mass spectrum of the precursor ion. **(b)** Sequence map of the phosphorylated peptide showing bond cleavages and phosphorylation sites. Cysteine carboxymethylation is considered in sequence mapping. **(c)** Representative fragment ions from ECD fragmentation. Fragment ions confirmed phosphorylation on Ser640, Ser645 and Ser652. Expt’l, experimental monoisotopic mass based on data obtained from MS or MS/MS experiments. Calc’d, calculated monoisotopic mass based on amino acid sequences.

**Figure S4.**
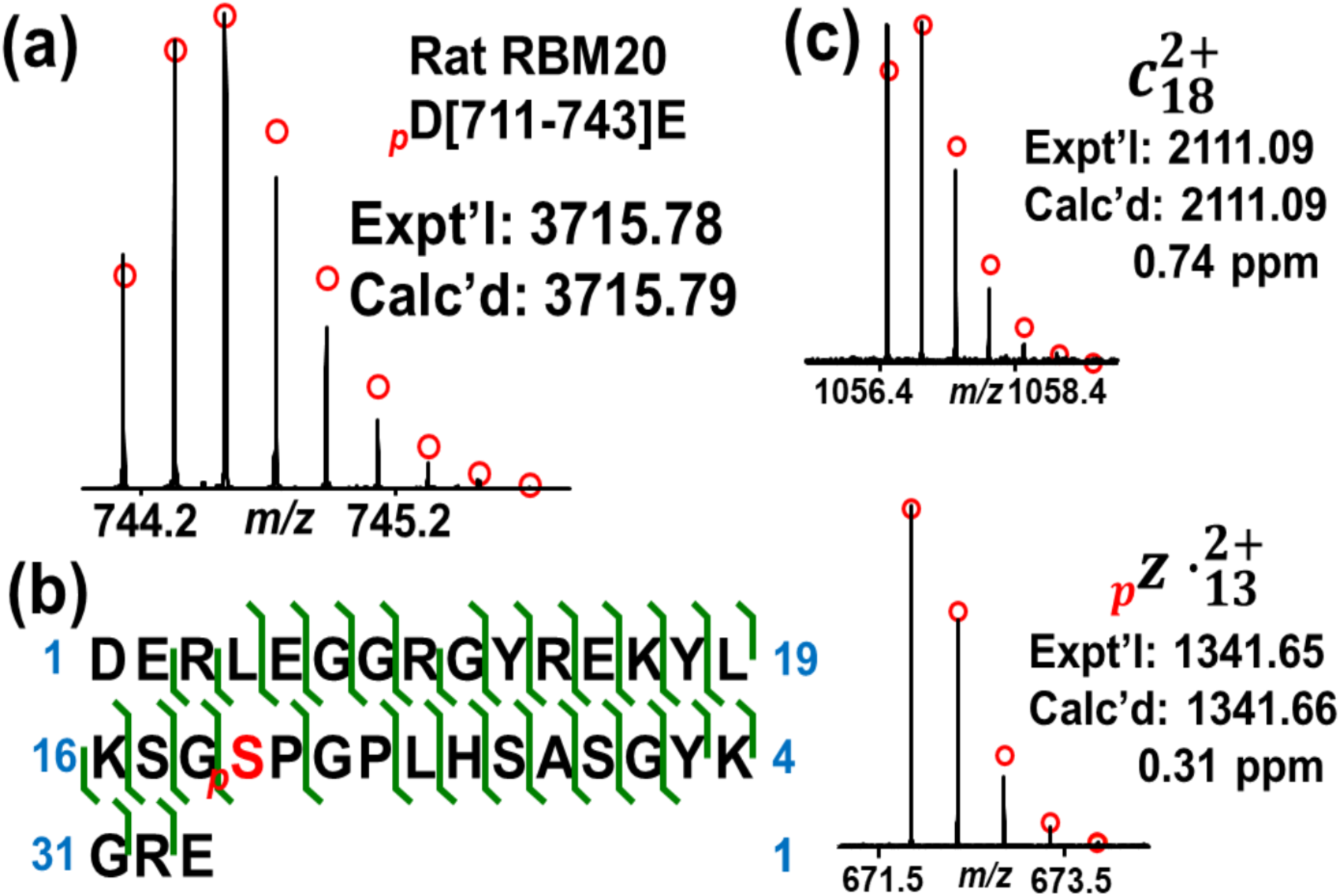
Identification of a phosphorylated peptide D[711-743]E with one phosphorylation site, Ser729, from Asp-N digestion of rat RBM20 (RBM20_Rat, UniProtKB-E9PT37). **(a)** Mass spectrum of the precursor ion. **(b)** Sequence map of the phosphorylated peptide showing bond cleavages and phosphorylation sites. Cysteine carboxymethylation is considered in sequence mapping. **(c)** Representative fragment ions from ECD fragmentation. Fragment ions confirmed phosphorylation on Ser729.

**Figure S5.**
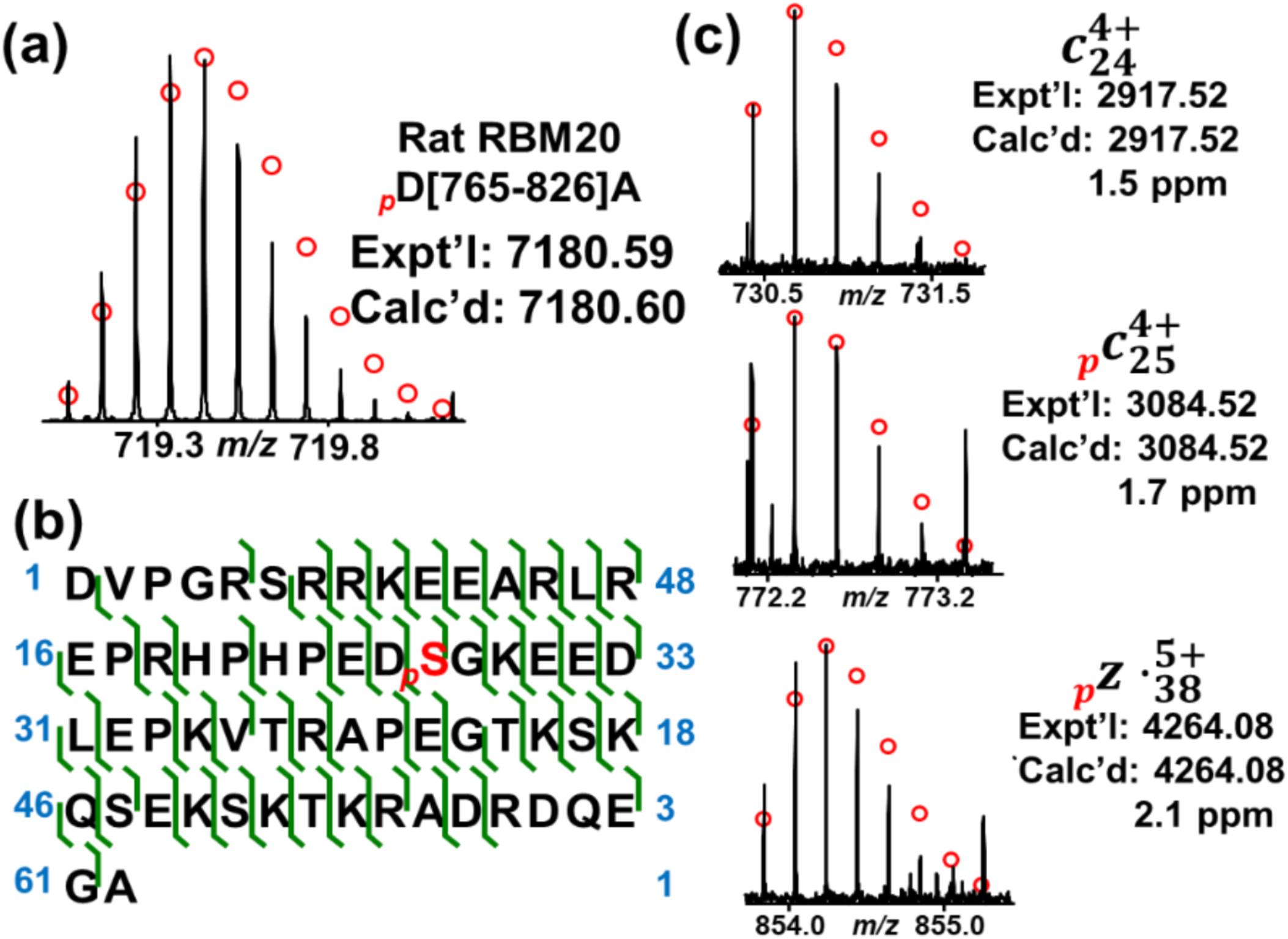
Identification of a phosphorylated peptide D[765-826]A with one phosphorylation site, Ser789, from Asp-N digestion of rat RBM20 (RBM20_Rat, UniProtKB-E9PT37). **(a)** Mass spectrum of the precursor ion. **(b)** Sequence map of the phosphorylated peptide showing bond cleavages and phosphorylation sites. Cysteine carboxymethylation is considered in sequence mapping. **(c)** Representative fragment ions from ECD fragmentation. Fragment ions confirmed phosphorylation on Ser789.

**Figure S6.**
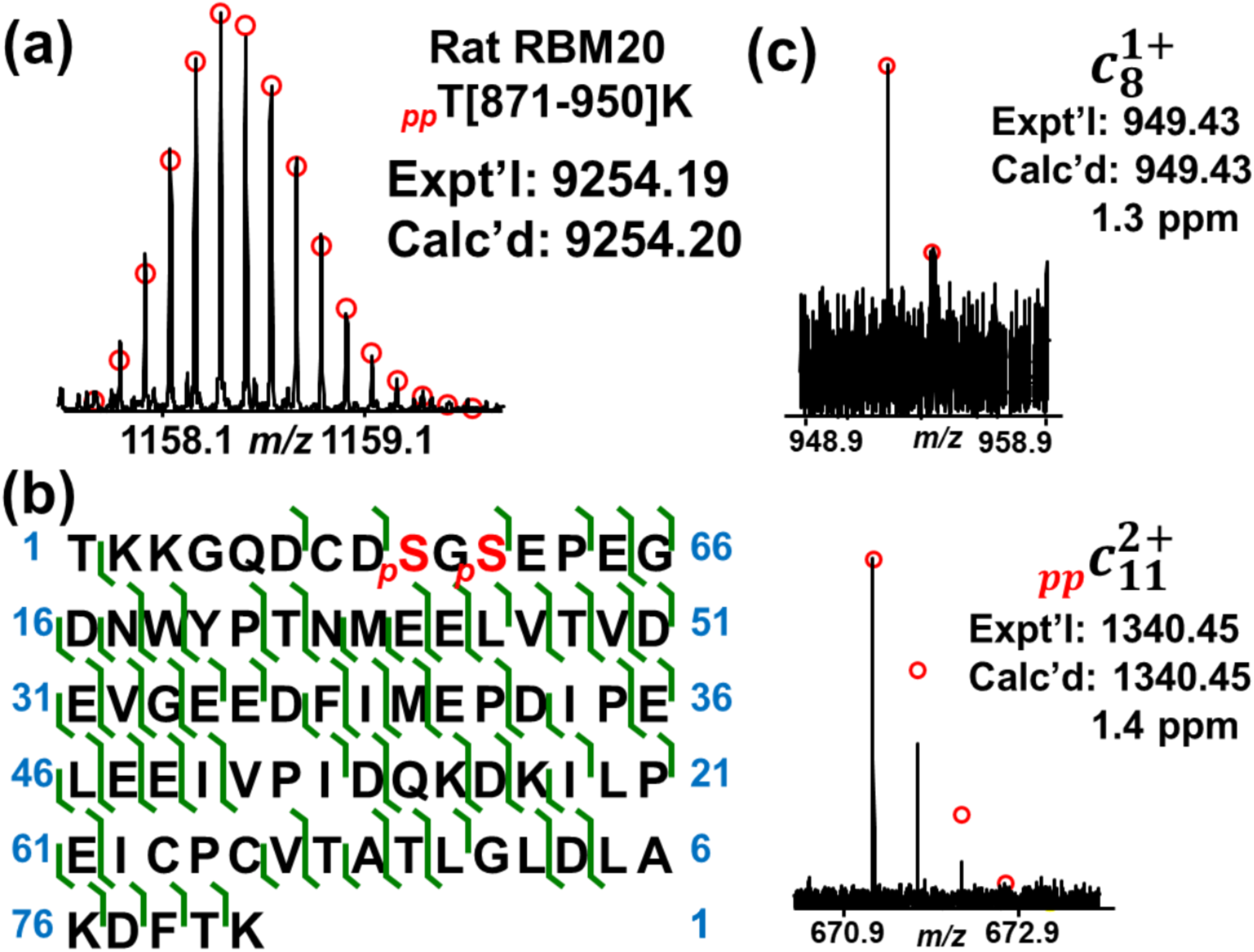
Identification of a phosphorylated peptide T[871-950]K with two phosphorylation sites, Ser879 and Ser881, from Trypsin digestion of rat RBM20 (RBM20_Rat, UniProtKB-E9PT37). **(a)** Mass spectrum of the precursor ion. **(b)** Sequence map of the phosphorylated peptide showing bond cleavages and phosphorylation sites. Cysteine carboxymethylation is considered in sequence mapping. **(c)** Representative fragment ions from ECD fragmentation. Fragment ions confirmed phosphorylation on Ser879 and Ser881.

**Figure S7.**
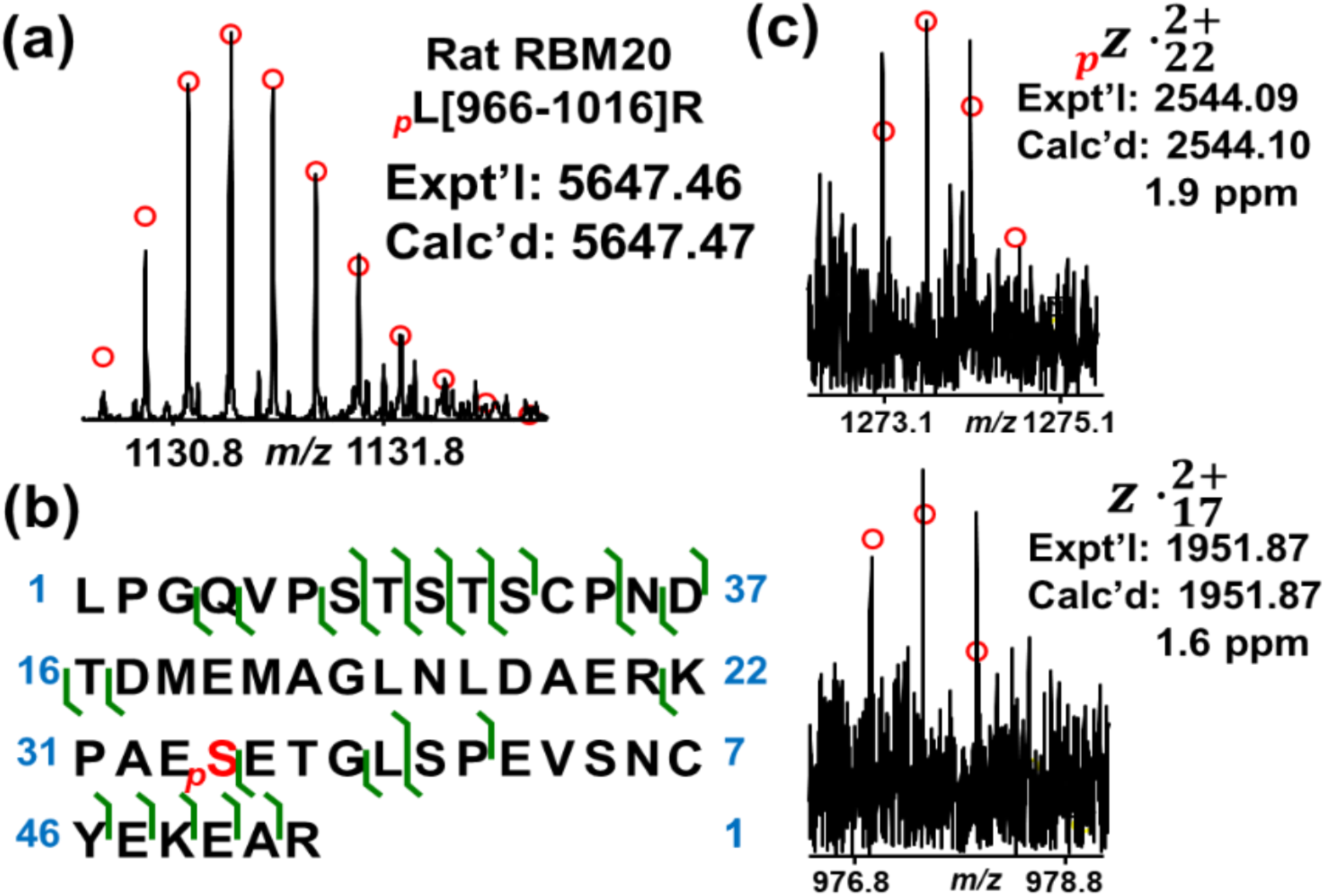
Identification of a phosphorylated peptide L[966-1016]R with one phosphorylation site, Ser999, from Trypsin digestion of rat RBM20 (RBM20_Rat, UniProtKB-E9PT37). **(a)** Mass spectrum of the precursor ion. **(b)** Sequence map of the phosphorylated peptide showing bond cleavages and phosphorylation sites. Cysteine carboxymethylation is considered in sequence mapping. **(c)** Representative fragment ions from ECD fragmentation. Fragment ions confirmed phosphorylation on Ser999.

**Figure S8.**
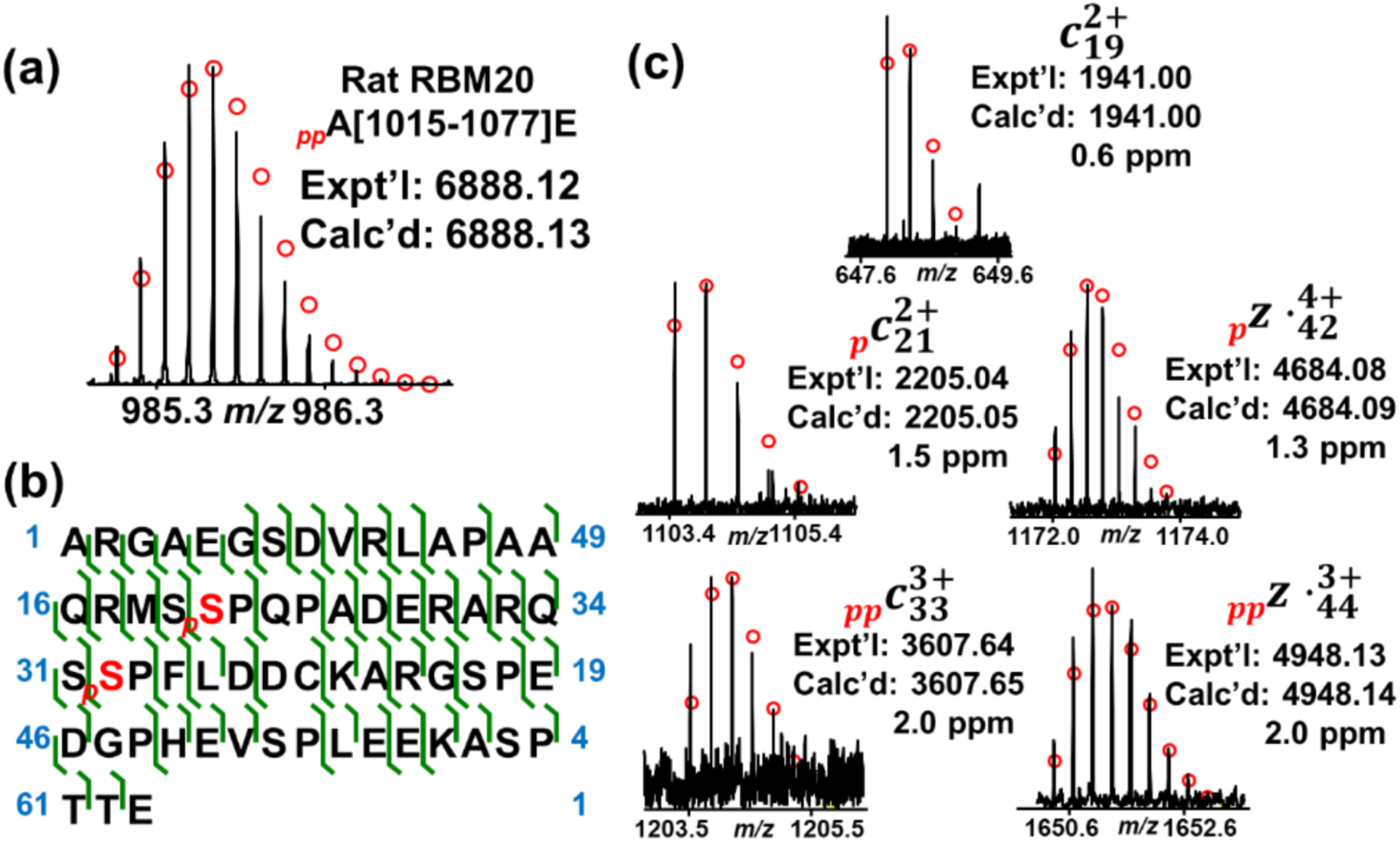
Identification of a phosphorylated peptide A[1015-1077]E with two phosphorylation sites, Ser1034 and Ser1046, from Glu-C digestion of rat RBM20 (RBM20_Rat, UniProtKB-E9PT37). **(a)** Mass spectrum of the precursor ion. **(b)** Sequence map of the phosphorylated peptide showing bond cleavages and phosphorylation sites. Cysteine carboxymethylation is considered in sequence mapping. **(c)** Representative fragment ions from ECD fragmentation. Fragment ions confirmed phosphorylation on Ser1034 and Ser1046.

**Figure S9.**
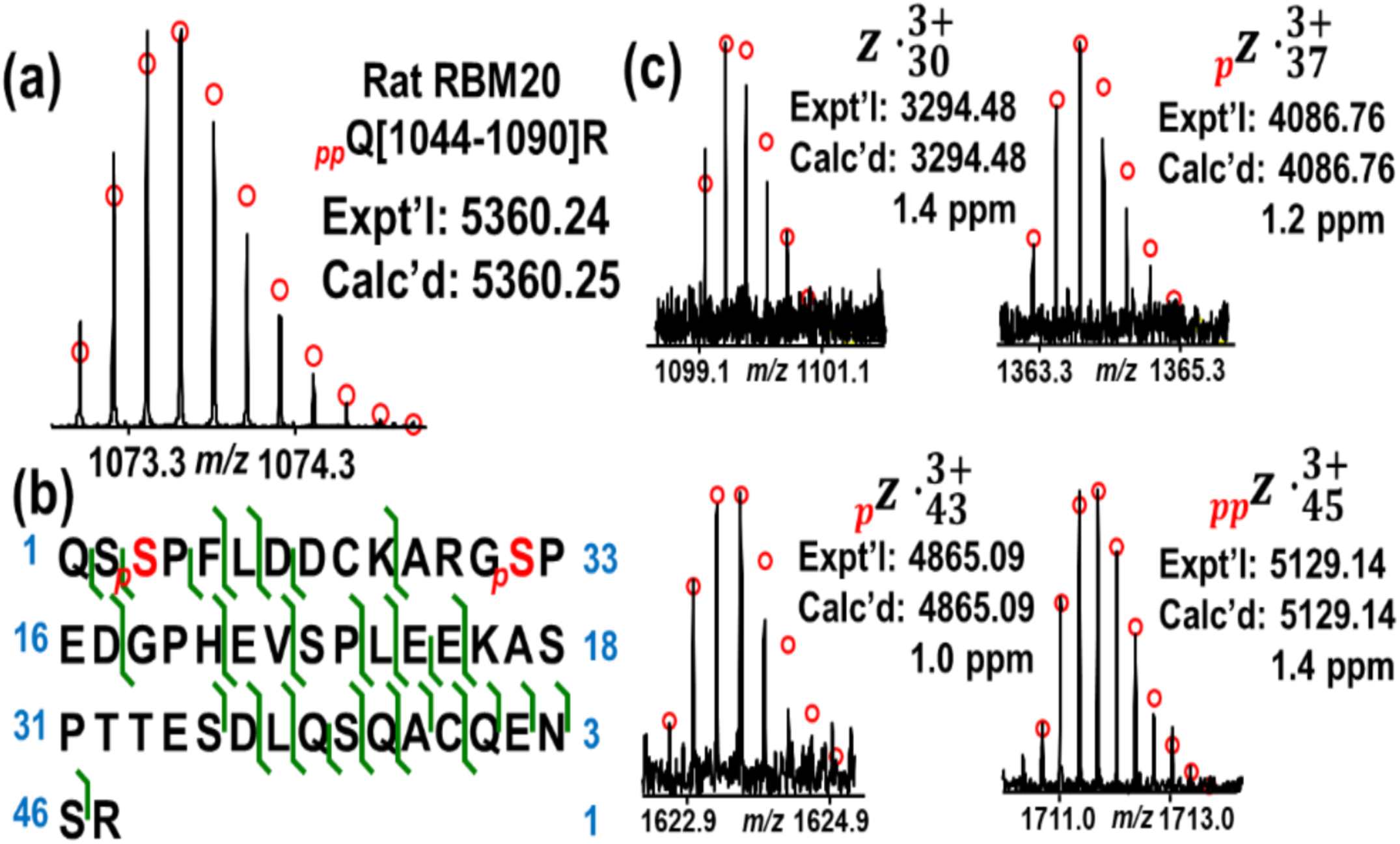
Identification of a phosphorylated peptide Q[1011-1090]R with two phosphorylation sites, Ser1046 and Ser1057, from Trypsin digestion of rat RBM20 (RBM20_Rat, UniProtKB-E9PT37). **(a)** Mass spectrum of the precursor ion. **(b)** Sequence map of the phosphorylated peptide showing bond cleavages and phosphorylation sites. Cysteine carboxymethylation is considered in sequence mapping. **(c)** Representative fragment ions from ECD fragmentation. Fragment ions confirmed phosphorylation on Ser1046 and Ser1057.

**Figure S10.**
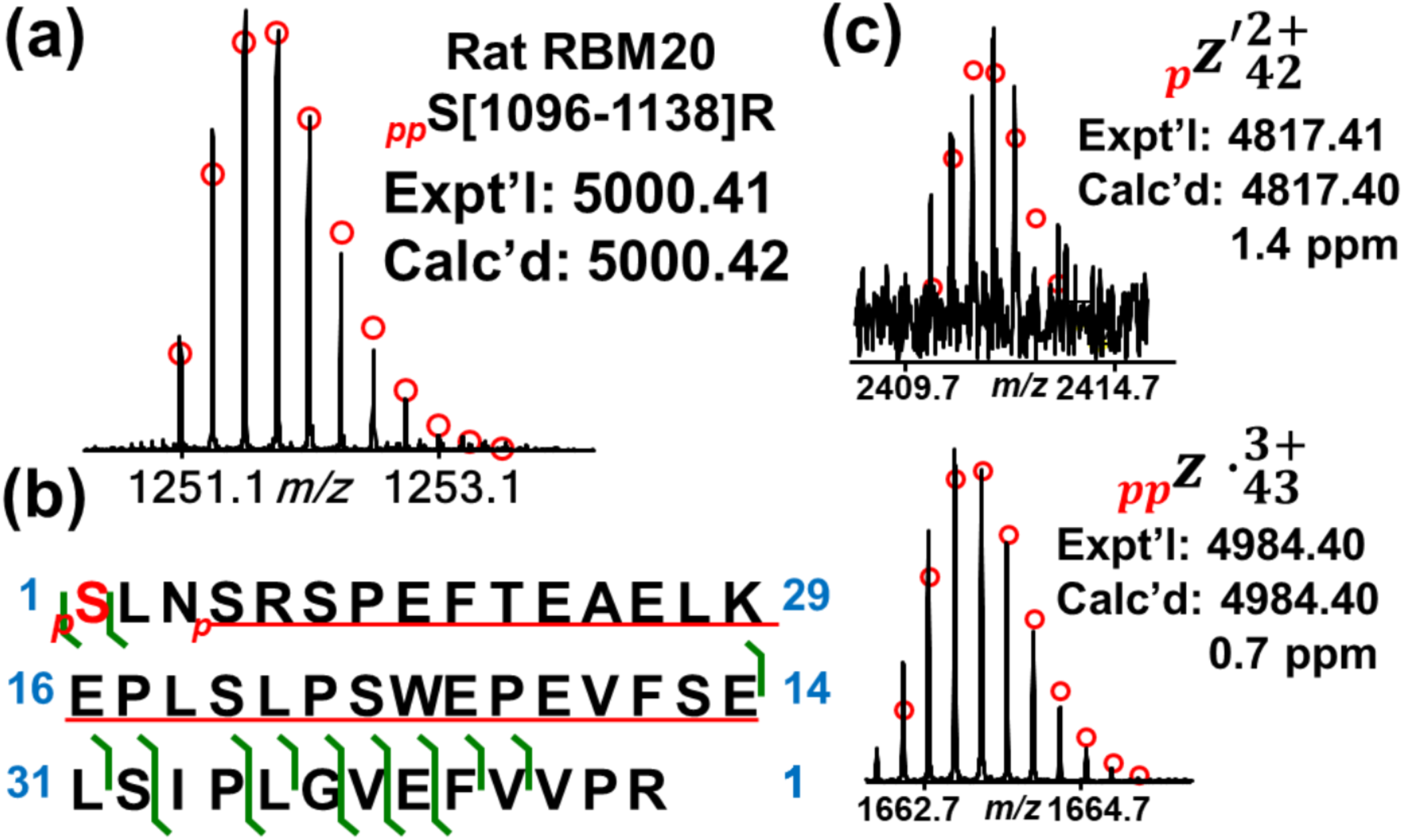
Identification of a phosphorylated peptide S[1096-1138]R with one phosphorylation site, Ser1096, from Trypsin digestion of rat RBM20 (RBM20_Rat, UniProtKB-E9PT37). **(a)** Mass spectrum of the precursor ion. **(b)** Sequence map of the phosphorylated peptide showing bond cleavages and phosphorylation sites. Cysteine carboxymethylation is considered in sequence mapping. The amino acids with underline show the range of another phosphorylation site which cannot be localized based on current MS/MS data. **(c)** Representative fragment ions from ECD fragmentation. Fragment ions confirmed phosphorylation on Ser1096.

**Figure S11.**
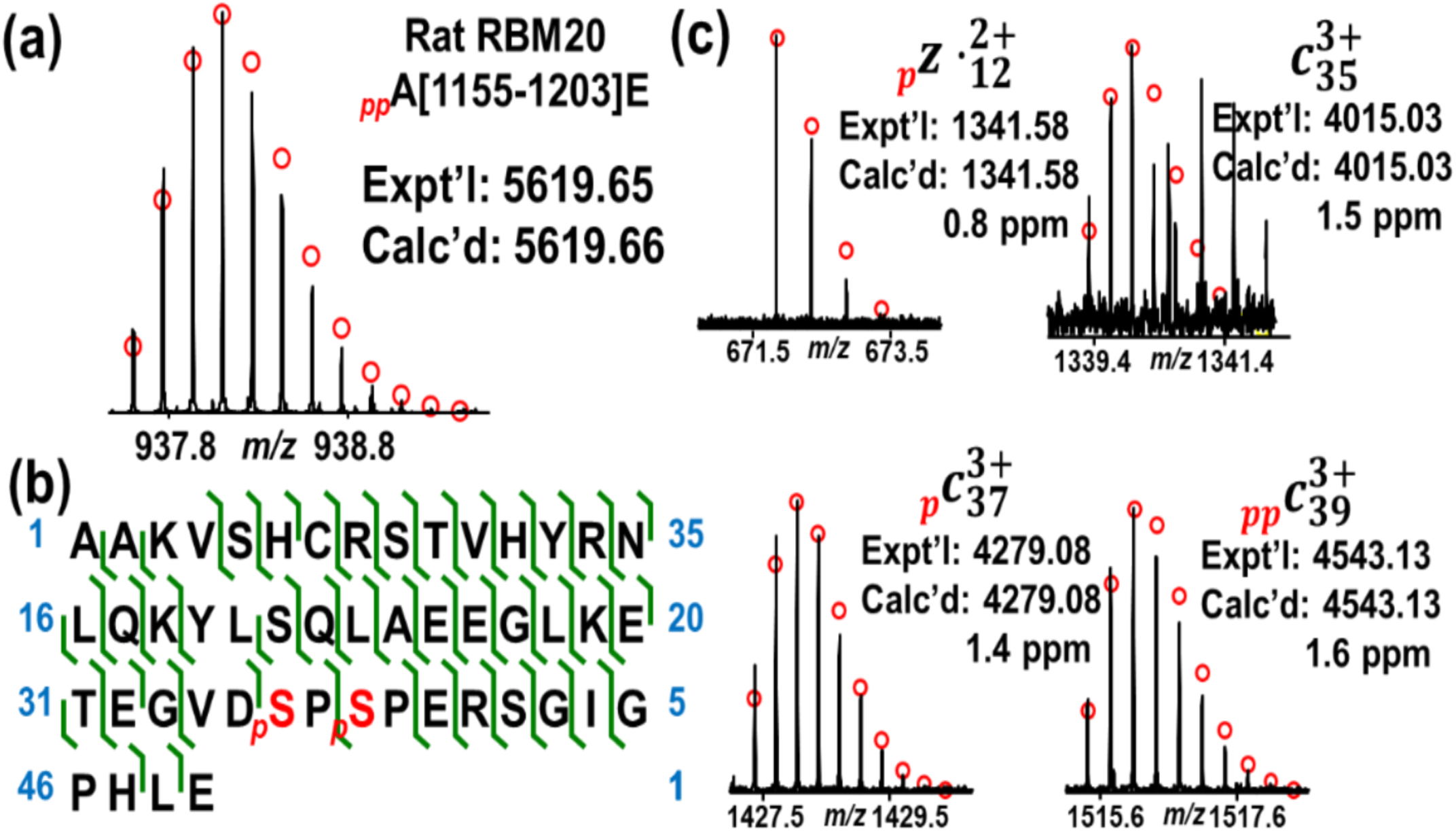
Identification of a phosphorylated peptide A[1155-1203]E with two phosphorylation sites, Ser1190 and Ser1192, from Glu-C digestion of rat RBM20 (RBM20_Rat, UniProtKB-E9PT37). **(a)** Mass spectrum of the precursor ion. **(b)** Sequence map of the phosphorylated peptide showing bond cleavages and phosphorylation sites. Cysteine carboxymethylation is considered in sequence mapping. **(c)** Representative fragment ions from ECD fragmentation. Fragment ions confirmed phosphorylation on Ser1190 and Ser1192

**Figure S12.**
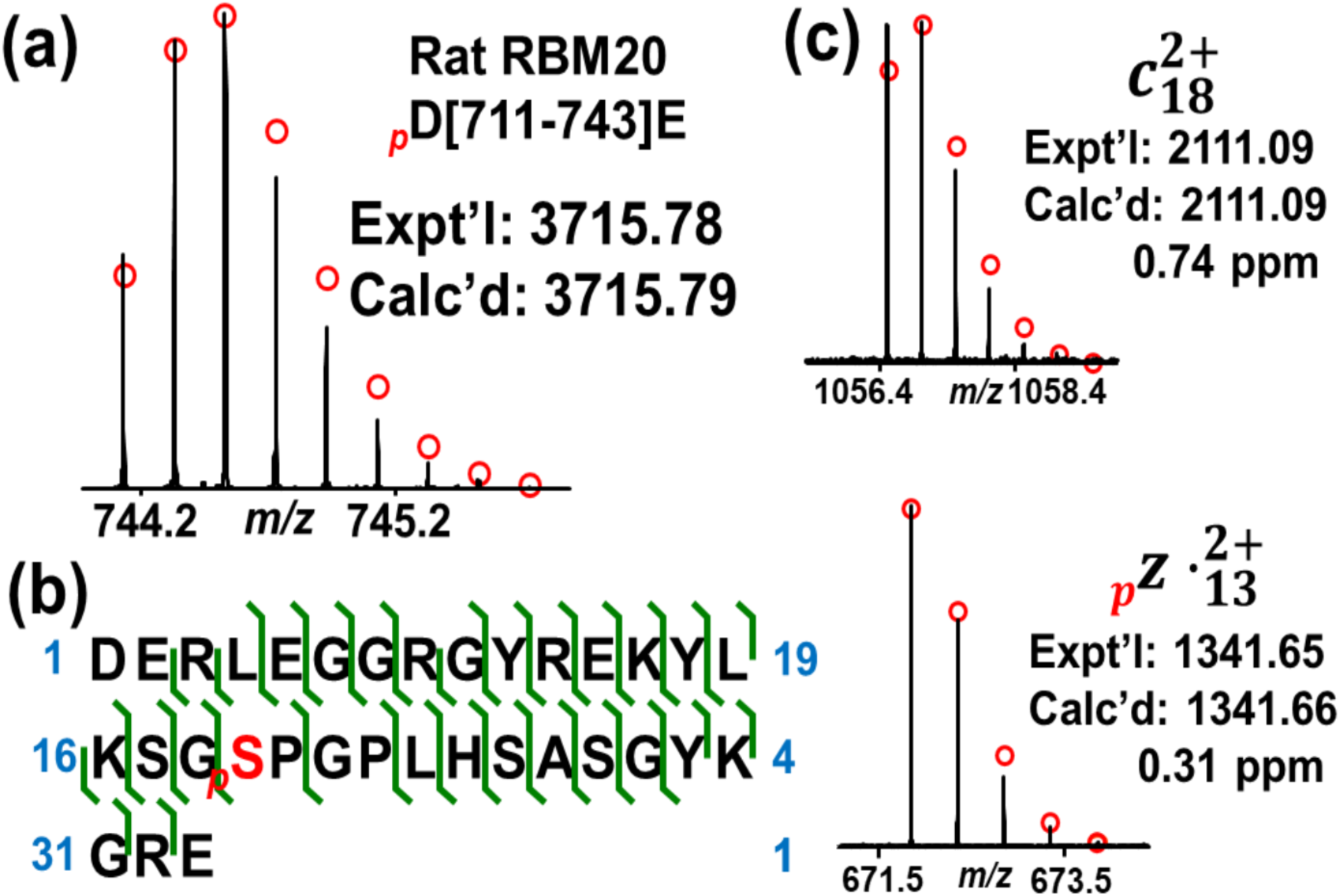
Summary of all identified RBM20 phosphorylation sites. Phosphorylated serine is marked in red. RS domain is highlighted in blue.

**Figure S13.**
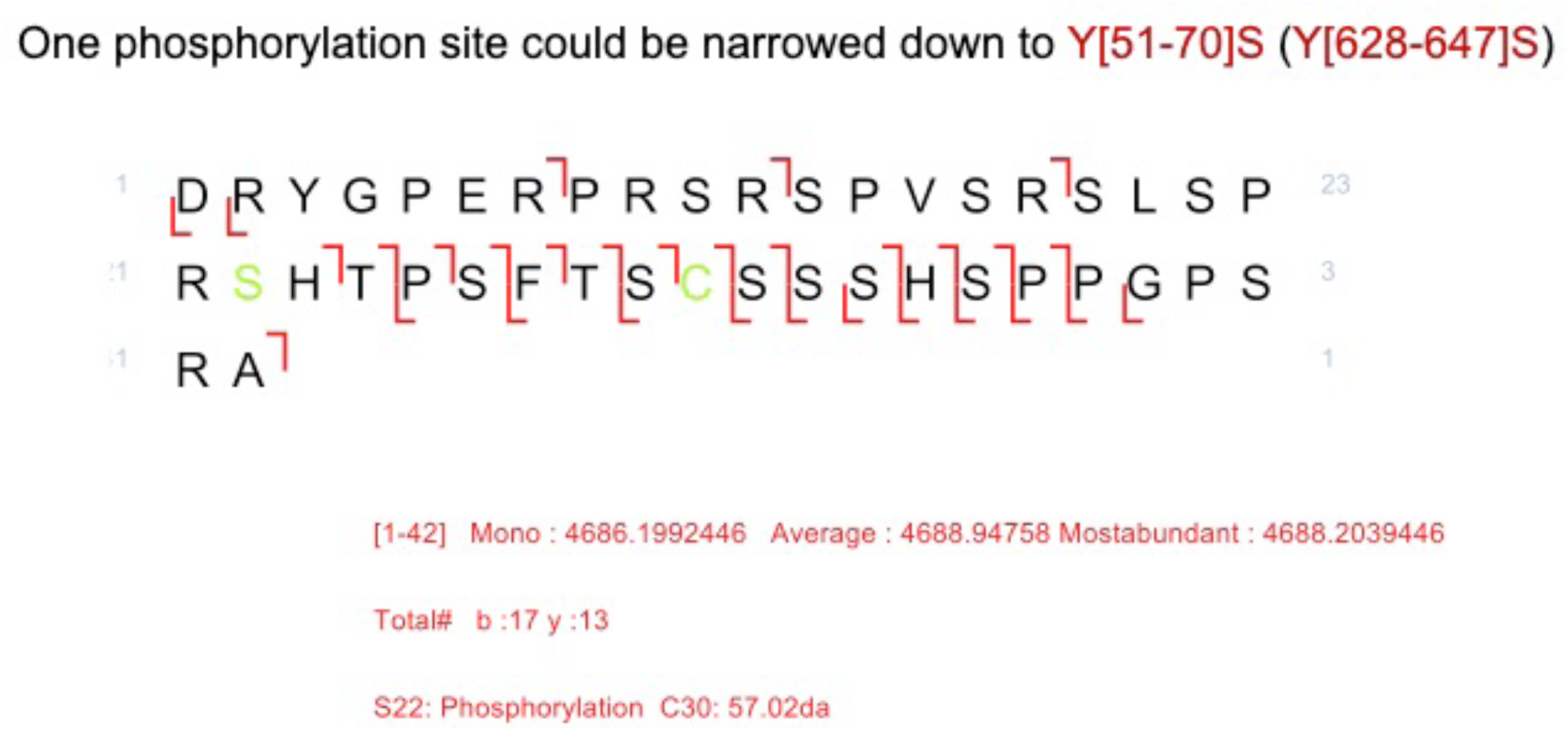
LC-MS/MS analysis of samples from the in vitro kinase assay showing the presence of phosphorylation in peptide D[626-667]A.

## References

1. Ziaeian B, and Fonarow GC. Epidemiology and aetiology of heart failure. Nat Rev Cardiol. 2016;6(13):368–378.

2. Hunt SA, Abraham WT, Chin MH, Feldman AM, Francis GS, and Ganiats TG, et al. Focused update incorporated into the ACC/AHA 2005 Guidelines for the Diagnosis and Management of Heart Failure in Adults A Report of the American College of Cardiology Foundation/American Heart Association Task Force on Practice Guidelines Developed in Collaboration With the International Society for Heart and Lung Transplantation. J Am Coll Cardiol. 2009; 53(15):e1–e90.

3. von Lueder TG, and Krum H. New medical therapies for heart failure. Nat Rev Cardiol. 2015; 12(12):730–740.

4. Greaser, M.L. et al. Developmental changes in rat cardiac titin/connectin: transitions in normal animals and in mutants with a delayed pattern of isoform transition. J. Muscle. Res. Cell. Motil. 26, 325–332 (2005).

5. Greaser, M.L. et al. Mutation that dramatically alters rat titin isoform expression and cardiomyocyte passive tension. J. Mol. Cell. Cardiol. 44, 983–991 (2008).

6. Guo, W. et al. RBM20, a gene for hereditary cardiomyopathy, regulates titin splicing. Nat. Med. 18, 766–773 (2012).

7. Brauch, K.M. et al. Mutations in ribonucleic acid binding protein gene cause familial dilated cardiomyopathy. J. Am. Coll. Cardiol. 54, 930–941 (2009).

8. Millat, G. et al. Clinical and mutational spectrum in a cohort of 105 unrelated patients with dilated cardiomyopathy. Eur. J. Med. Genet. 54, e570–e575 (2011).

9. Monaco, I. et al. Double de novo mutations in dilated cardiomyopathy with cardiac arrest. J. Electrocardiol. 53, 40–43 (2019).

10. Pantou, M.P. et al. Phenotypic Heterogeneity within members of a family carrying the same RBM20 mutation R634W. Cardiology. 141, 150–155 (2018).

11. Refaat, M.M. et al. Genetic variation in the alternative splicing regulator RBM20 is associated with dilated cardiomyopathy. Heart. Rhythm. 9, 390–396 (2012).

12. Rampersaud, E. et al. Rare variant mutations identified in pediatric patients with dilated cardiomyopathy. Prog. Pediatr. Cardiol. 31, 39–47 (2011).

13. Li, D. et al. Identification of novel mutations in RBM20 in patients with dilated cardiomyopathy. Clin. Transl. Sci. 3, 90–97 (2010).

14. Guo, W., Pleitner, J.M., Saupe, K.W. & Greaser, M.L. Pathophysiological defects and transcriptional profiling in the RBM20^-/-^ rat model. PLOS. One. 8, e84281 (2013).

15. Li, S., Guo, W., Dewey, C.N. & Greaser, M.L. Rbm20 regulates titin alternative splicing as a splicing repressor. Nucleic. Acids. Res. 41, 2659–2672 (2013).

16. Methawasin, M. et al. Experimentally increasing titin compliance in a novel mouse model attenuates the Frank-Starling mechanism but has a beneficial effect on diastole. Circulation. 129, 1924–1936 (2014).

17. Maatz, H. et al. RNA-binding protein RBM20 represses splicing to orchestrate cardiac pre-mRNA processing. J. Clin. Invest. 124, 3419–3430 (2014).

18. van den Hoogenhof, M. et al. RBM20 mutations induce an arrhythmogenic dilated cardiomyopathy related to disturbed calcium handling. Circulation. 138, 1330–1342 (2018).

19. Guo, W. et al. Splicing factor RBM20 regulates transcriptional network of Titin associated and calcium handling genes in the heart. Int J Biol Sci. 14, 369–380 (2018).

20. Serrano, P. et al. Directional phosphorylation and nuclear transport of the splicing factor SRSF1 is regulated by an RNA recognition motif. J. Mol. Biol. 428, 2430–2445 (2016).

21. Zhu, C., Chen, Z. & Guo, W. Pre-mRNA mis-splicing of sarcomeric genes in heart failure. Biochim. Biophys. Acta. Mol. Basis. Dis. 1863, 2056–2063 (2017).

22. Rexiati, M., Sun, M. & Guo, W. Muscle-specific mis-splicing and heart disease exemplified by RBM20. Genes (Basel). 9, 18–31 (2018).

23. Ghosh, G. & Adams, J.A. Phosphorylation mechanism and structure of serine-arginine protein kinases. FEBS J. 278, 587–597 (2011).

24. Corkery, D.P., Holly, A.C., Lahsaee, S. & Dellaire, G. Connecting the speckles: Splicing kinases and their role in tumorigenesis and treatment response. Nucleus-Phila. 6, 279–288 (2015).

25. Zhou, Z. & Fu, X.D. Regulation of splicing by SR proteins and SR protein-specific kinases. Chromosoma. 122, 191–207 (2013).

26. Naro, C. & Sette, C. Phosphorylation-mediated regulation of alternative splicing in cancer. Int.J. Cell. Biol. 2013, 151839 (2013).

27. Prasad, J., Colwill, K., Pawson, T. & Manley, J.L. The protein kinase Clk/Sty directly modulates SR protein activity: both hyper-and hypophosphorylation inhibit splicing. Mol. Cell. Biol. 19, 6991–7000 (1999).

28. Lai, M.C., Lin, R.I., Huang, S.Y., Tsai, C.W. & Tarn, W.Y. A human importin-beta family protein, transportin-SR2, interacts with the phosphorylated RS domain of SR proteins. J. Biol. Chem. 275, 7950–7957 (2000).

29. Lai, M.C., Lin, R.I. & Tarn, W.Y. Transportin-SR2 mediates nuclear import of phosphorylated SR proteins. Proc. Natl. Acad. Sci. USA. 98, 10154–10159 (2001).

30. Mermoud, J.E., Cohen, P.T. & Lamond, A.I. Regulation of mammalian spliceosome assembly by a protein phosphorylation mechanism. EMBO J. 13, 5679–5688 (1994).

31. Lai, M.C. & Tarn, W.Y. Hypophosphorylated ASF/SF2 binds TAP and is present in messenger ribonucleoproteins. J. Biol. Chem. 279, 31745–31749 (2004).

32. Jin, Y. et al. Complete characterization of cardiac myosin heavy chain (223 kDa) enabled by size-exclusion chromatography and middle-down mass spectrometry. Anal. Chem. 89, 4922–4930 (2017).

33. Chen, B. et al. Middle-down multi-attribute analysis of antibody-drug conjugates with electron transfer dissociation. Anal. Chem. 18, 11661–11669 (2019).

34. Zhu, C. & Guo, W. Detection and quantification of the giant protein titin by SDS-agarose gel electrophoresis. MethodsX. 4, 320–327 (2017).

35. Zhu, C. et al. RBM20 is an essential factor for thyroid hormone-regulated titin isoform transition. J. Mol. Cell. Biol. 7, 88–90 (2015).

36. Zhu, C., Yin, Z., Tan, B. & Guo, W. Insulin regulates titin pre-mRNA splicing through the PI3K-Akt-mTOR kinase axis in a RBM20-dependent manner. Biochim Biophys Acta. Mol. Basis. Dis. 1863, 2363–2371 (2017).

37. Cai, W. et al. Unbiased proteomics method to assess the maturation of human pluripotent stem cell-derived cardiomyocytes. Circ. Res. 125, 936–953 (2019).

38. Lin, Z. et al. Simultaneous quantification of protein expression and modifications by top-down targeted proteomics: a case of sarcomeric subproteome. Mol. Cell. Proteomics. 18, 594–605 (2019).

39. Jin, Y. et al. Comprehensive characterization of monoclonal antibody by Fourier transform ion cyclotron resonance mass spectrometry. MAbs. 11, 106–115 (2019).

40. Liu, X. et al. Identification of ultramodified proteins using top-down tandem mass spectra. J. Proteome. Res. 12, 5830–5838 (2013).

41. Guner, H. et al. MASH Suite: a user-friendly and versatile software interface for high-resolution mass spectrometry data interpretation and visualization. J. Am. Soc. Mass. Spectrom. 25, 464–470 (2014).

42. Cai, W. et al. MASH Suite Pro: A comprehensive software tool for Top-down proteomics. Mol. Cell. Proteomics. 15, 703–714 (2016).

43. Polymenidou, M. et al. Long pre-mRNA depletion and RNA missplicing contribute to neuronal vulnerability from loss of TDP-43. Nat. Neurosci. 14, 459–468 (2011).

44. Vanderweyde, T., Youmans, K., Liu-Yesucevitz, L. & Wolozin, B. Role of stress granules and RNA-binding proteins in neurodegeneration: a mini-review. Gerontology. 59, 524–533 (2013).

45. Buratti, E. & Baralle, F.E. Multiple roles of TDP-43 in gene expression, splicing regulation, and human disease. Front. Biosci. 13, 867–878 (2008).

46. Kuo, P.H., Doudeva, L.G., Wang, Y.T., Shen, C.K. & Yuan, H.S. Structural insights into TDP-43 in nucleic-acid binding and domain interactions. Nucleic. Acids. Res. 37, 1799–1808 (2009).

47. Murayama, R. et al. Phosphorylation of the RSRSP stretch is critical for splicing regulation by RNA-Binding Motif Protein 20 (RBM20) through nuclear localization. Sci. Rep. 8, 1–14 (2018).

48. Lin, S. & Fu, X. SR proteins and related factors in alternative splicing. Adv. Exp. Med. Biol. 623, 107–122 (2007).

49. Lukong, K.E., Chang, K.W., Khandjian, E.W. & Richard, S. RNA-binding proteins in human genetic disease. Trends. Genet. 24, 416–425 (2008).

50. Wang, G.S. & Cooper, T.A. Splicing in disease: disruption of the splicing code and the decoding machinery. Nat. Rev. Genet. 8, 749–761 (2007).

51. Cooper, T.A., Wan, L. & Dreyfuss, G. RNA and disease. Cell. 136, 777–793 (2009).

52. Cáceres, J.F., Screaton, G.R. & Krainer, A.R. A specific subset of SR proteins shuttles continuously between the nucleus and the cytoplasm. Genes. Dev. 12 (1):55–66 (1998).

53. Piñol-Roma, S. & Dreyfuss, G. Shuttling of pre-mRNA binding proteins between nucleus and cytoplasm. Nature. 355, 730–732 (1992).

54. Gabut, M., Chaudhry, S. & Blencowe, B.J. SnapShot: The splicing regulatory machinery. Cell. 133, 191–192 (2008).

55. Daoud, H. et al. Contribution of TARDBP mutations to sporadic amyotrophic lateral sclerosis. J. Med. Genet. 46, 112–114 (2009.

56. Kabashi, E. et al. TARDBP mutations in individuals with sporadic and familial amyotrophic lateral sclerosis. Nat. Genet. 40, 572–574 (2008).

57. Rutherford, N.J. et al. Novel mutations in TARDBP (TDP-43) in patients with familial amyotrophic lateral sclerosis. PLOS. Genet. 4, e1000193 (2008).

58. Sreedharan, J. et al. TDP-43 mutations in familial and sporadic amyotrophic lateral sclerosis. Science. 319, 1668–1672 (2008).

59. Schneider, J. et al. A ribonucleoprotein-granule pathway to heart failure in human RBM20 cardiomyopathy gene-edited pigs. Nat. Med. (2020). Accepted.

60. Cai, H. et al. Angiotensin II influences Pre-mRNA splicing regulation by enhancing RBM20 transcription through activation of the MAPK/ELK1 signaling pathway. Int. J. Mol. Sci. 20, E5090 (2019).

